# Spatial Transcriptomics Reveal Developmental Dynamics of the Human Cerebral Cortex and Striatum

**DOI:** 10.1101/2025.08.14.670213

**Authors:** Yunjia Zhang, Youning Lin, Xunan Shen, Xue Xiao, Cai Song, Xiaobo Shi, Tao Zhou, Yanxin Li, Zihan Wu, Zhenkun Zhuang, Chunqiong Li, Meng Li, Feng Wen, Jianlin Liu, Qiangqiang Zhang, Zhao-Lu Li, Songbo Zhang, Lei Cao, Susu Qu, Yaqi Li, Jianhua Yao, Fubaoqian Huang, Xin Liu, Ziqing Deng, Longqi Liu, Xun Xu, Jianwei Jiao, Li Zhang, Shiping Liu, Yaoyao Zhang

## Abstract

The human fetal brain undergoes morphological changes that contribute to the development of regional functionalities. However, the features of structural development, the underlying molecular and cellular signatures in the fetal brain remain unclear. With spatial transcriptomics and snRNA-seq, we identified 25 forebrain regions and characterized the dynamic changes in the cortex and striatum during the late first and early second trimesters. In particular, we discovered that temporal lobe enriched NPY-expressing L2/3 EX neuron potentially interacted with L4 EX neurons during cortical expansion and arealization. Additionally, the gyrus and sulcus were developmental asynchronous, in which *HOPX* and *SPARC* genes were potentially involved. Further investigation on the striatum showed specific genes and cell types that enriched in patch and matrix compartments, and *SST*-positive interneurons potentially involved in the development of these structures. Together, our results give insights into the understanding of early fetal brain development.

## Introduction

The fetal brain undergoes dynamic morphological and cellular alterations during development, resulting in a mature configuration of the adult brain, especially for the most evolved neocortex with convoluted, layered gray matter as well as the striatum related to cognitive control function^1,2^.

During corticogenesis, neurons assemble in an inside-out layering manner. The laminar structure expands and undergoes transient changes, accompanied by fine-tuned layering and cortical arealization^3^. However, the precise spatiotemporal distribution of cell types across cortical regions during development at single_cell resolution hasn’t been resolved yet. Furthermore, whether distinct neuronal subtypes emerge in a region-specific manner during corticogenesis is unknown. Another prominent and evolutionary feature of the human brain is the folding of the neocortex^4^. The complex folding structure, which can be measured as a high gyrification index^5^, provides the possibility for a correlation with human-specific lobes. Diverse cellular and genetic mechanisms contribute to the folding structures. However, how convoluted structures systematically form in the developing human brain is unclear. These mentioned several unknowns in the cortex were the aims of the current study.

The cortex is widely connected with the basal ganglia via segregated populations of striatal projection neurons^6^. Over 90% of striatal projection neurons are striatal medium spiny neurons (MSNs)^7^. Previous studies have shown the development of human MSNs in the proliferating zone of the lateral ganglionic eminence (LGE), where decoded TFs and lincRNAs regulate the differentiation of preMSNs into D1 and D2 MSNs^8^. However, the development and distribution of D1/D2 hybrid MSNs remain unknown. In the mantle zone, MSNs and interneurons develop into a nucleus consisting of patch and matrix compartments in the basal ganglionic system. Studies on radiolabeled postmitotic neurons in rat embryos showed that patch neurons were born earlier than matrix neurons, and therefore established connections with the substantia nigra earlier^9, 10^. While to date, the cellular properties involved in the development of patches and matrices, as well as the molecular dynamics driving structural identities are unclear. Thus, these two unclears in the striatum are the other aims of the current study.

To address these questions on nonpathological human fetal tissue, a combination of spatiotemporal seq and snRNA seq was used in the present study, which provides an opportunity to understand the structural development and changes of molecular and cellular signatures in the cortex and striatum. This resource is interactively accessible at: http://db.cngb.org/cnsa/project/CNP0005272_8c0dc42f/reviewlink/

## Results

### Spatially resolved cellular map of the human fetal brain

To comprehensively characterize the dynamics of spatiotemporal gene expression dynamics in developing human fetal brains, we employed an integrated multi-omics approach combining high-resolution spatial transcriptomics (stereo-seq^11^) and single-nucleus RNA sequencing (snRNA-seq; DNBelab C4^11^). The spatial profile was conducted on coronal sections across cortical and striatal regions at gestational week (GW) 11,15, 21, and included data from Li et.al^12^, resulting in spatial data spanning GW8 to GW21, and encompassing first trimester (GW8, 10-12), and second trimester (GW15-16, 21) developmental stages. Single cell resolution was achieved by a cell segmentation algorithm^13^. Parallel snRNA-seq analysis was performed on histologically matched anatomical sections (coronal) to resolve cell-type-specific transcriptional signatures at corresponding developmental timepoints (**Fig 1A**). A total of 424,527 cells were generated by snRNA-seq. After integrated the published data^12, 14, 15^, we finally approached 798,138 cells for the human fetal brain. All the cells were clustered into 12 major cell types and 136 subtypes (**Figs 1B**, **S1A-D**).

The spatial brain regions in each coronal section were delineated using FuseMap^16^ at single cell resolution, as well as SpaGCN clustering^17^ at bin100 resolution, leading to an identification of 25 subregions within the developing brain, which were confirmed through the canonical marker genes (**Figs 1C, 1D, S2A-F, S3A, S3B**). The cortex was consisted of the ventricle zone (VZ) (*VIM, SOX2*)^18^, subventricular zone (SVZ) (*PPP1R17, EOMES*)^19^, intermediate zone (IZ, identified based on anatomy structure), subplate (SP) (*ST18, NR4A2*)^20^ and cortical plate (CP) (*NEUROD2, TBR1*)^19^ (**Figs 1C, 1D, S2B, S2C, S2F**). The subpallial regions developed the hippocampus (HIP) (*NEUROD1, NR3C2)*^21, 22^, GE-proliferating zone (*GAD2*)^15^, LGE-proliferating zone (*PAX6, SIX3*)^15^, LGE-mantle zone (also promising striatum, STR) (*TAC1, EBF1*)^15^, MGE-proliferating zone (*NKX2-1*)^15^, CGE-proliferating zone (*NR2F2*)^15^, globus pallidus (GP) (*LHX8, LHX6*)^23^, amygdala (AMY) (*NRP2, NR2F2*)^24^, septum (SEP) (*ZIC4, ZIC1, ZIC2*)^25, 26^, claustrum (Cla) (*NTNG2, NR4A2*)^27, 28^, thalamus (THT) (*TCF7L2, LHX9*)^29^, hypothalamus (HYT) (*LHX5-AS1*)^30^, bed nucleus of striatal terminal (BNST) (C*CK, NTS, SST, ADRA1B*)^31^ and etc. (**Figs 1C, 1D, S2B, S2C, S2F, S3B**). Brain regions in sagittal sections from Li et.al were spatially clustered with SpaGCN at bin100 **(Figs S3C-E**), followed by single-cell resolution annotation (**Figs S4A, S4B**). The spatially resolved regions within the sagittal plane exhibited significant spatial congruence with established anatomical boundaries **(Fig S4C).** Both coronal and sagittal brain regions exhibited strong correlation across different developmental stages (**Fig S4D**). By single-cell data integration with spatial transcriptomic mapping, we identified the cerebral cortex as the regions with the most cell number across the sampled regions (**Fig 1E**).

**Fig.1.**
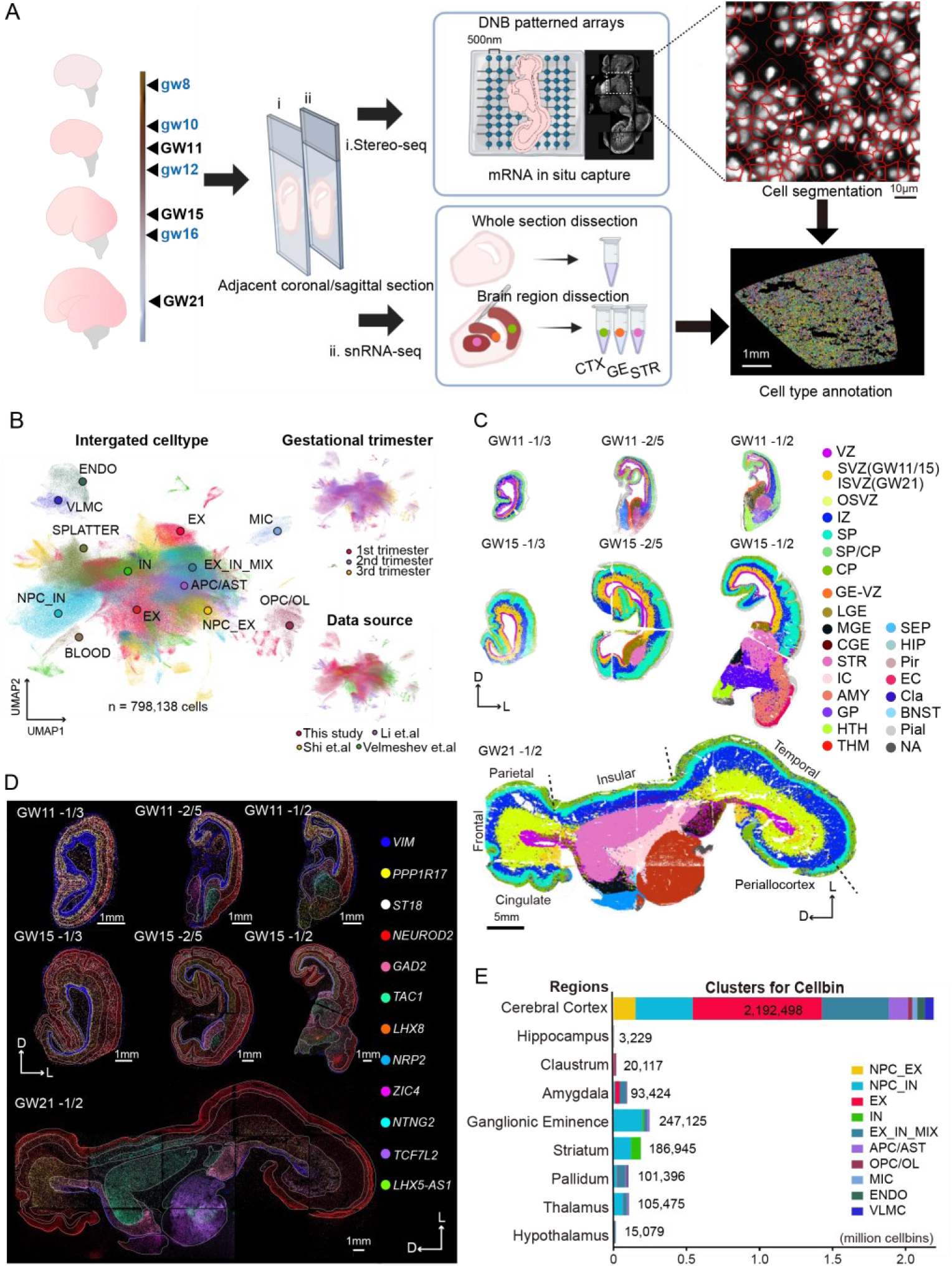
| Single-nuclei transcriptomics and stereo-seq characterize human fetal brain cellular map. **A.** Overall experimental design. Scales: 10μm (top); 1mm (bottom). GW: samples from this study; gw: published data by Li et al. **B.** UMAP showing snRNA-seq clustering and cell-type taxonomy. Cell types colored by the annotations (Left). EX: excitatory neuron; NPC_EX: neural progenitor cell_excitatory neuron; NPC_IN: neural progenitor cell_interneuron; IN: interneuron; EX_IN_MIX: excitatory neuron and interneuron mixed cells; APC/AST: astrocyte precursor cells/astrocyte; OPC/OL: oligodendrocyte precursor cells/oligodendrocyte; MIC: microglia; ENDO: endothelial cells; VLMC: vascular leptomeningeal cells. Cells colored by time points and data source (Right). **C.**Spatial clustering showing stereo-seq brain regions by FuseMap at single cell resolution. Brain regions are colored by the annotations. VZ: ventricle zone; SVZ: sub-ventricle zone; ISVZ: inner sub-ventricle zone; OSVZ: outer sub-ventricle zone; IZ: intermediate zone; SP: subplate; CP: cortical plate; LGE: lateral ganglionic eminence; MGE: medial ganglionic eminence; CGE: caudal ganglionic eminence; STR: striatum (caudate and putamen); IC: internal capsule; AMY: amygdala; GP: globus pallidus; HTH: hypothalamus; THM: thalamus; SEP: septum; HIP: hippocampus; Pir: piriform cortex; EC: entorhinal cortex; Cla: claustrum; BNST: bed nucleus of striata terminal; NA: not available. D: dorsal; L: lateral. Scales: 5mm. **D.**Spatial visualization showing specific brain region markers. White solid lines represent the borders of subregions. VZ: *VIM*, SVZ: *PPP1R17*, SP: *ST18*, CP: *NEUROD2*, GE-proliferating zone: *GAD2*, GP: *LHX8*, AMY: *NRP2*, SEP: *ZIC4*, Cla: *NTNG2*, THM: *TCF7L2*, HTH: *LHX5-AS1*. D: dorsal; L: lateral. Scales: 1mm. **E.**Stack-bar plot showing the distribution of clusters in brain regions. See also Figs. S1, S2, S3, S4.

### The dynamic distribution of excitatory cells in cortical expansion and arealization

Spatial clustering analyses demonstrated that the cortical plate developed above the subplate from GW11 to gw16 (**Fig 1C, S2C, S3C, S4A**), aligning with the prevailing theory of prelate division into the subplate and cortical plate around PCW12 (equivalent to GW14)^32^. To quantitatively analyze spatiotemporal cellular dynamics during neocortical expansion, stratified proportions of cells within embryonic cortical laminae were quantified across neurodevelopmental stages (**Fig 2A**). The VZ and SVZ were mainly occupied by the NPC_EX, NPC_IN and APC/AST (**Fig 2A, S5A**). In the NPC_EX cluster, the NPC_EX16 primarily localized in the VZ, exhibiting RG characteristics with *SOX2* expression, and NPC_EX2/7/1/6 predominantly resided in the SVZ and IZ, displaying IPC features with *PPP1R17* expression. In contrast, no distinct RG- or IPC-only clusters were identified within the NPC_IN population, the NPC_IN17/14/15/16/7 expressed both *SOX2* and *ASCL1*, initially localized to the VZ during first trimester (GW8–12) and expanded into SVZ during the second trimester (GW15–21); while NPC_IN6/12/11 distributed through SVZ, IZ and the cortical plate across the neurodevelopmental stages (**Figs 2A, S5B, S5C**).

A dynamic spatiotemporal redistribution of cellular populations was observed within the SP and CP laminae across progressive stages of corticogenesis. The distribution of excitatory neurons segregated into three distinct populations: EX26/19/45/10 localized in the SP by GW12, co-expressing *TLE4* and *BCL11B*, consistent with a deep layer neuron identity; EX5/29/30/2/11, early-born cortical neurons occupying SP/CP during the first trimester (GW8–12) and restricted to CP by the second trimester (GW15–21), highly expressing layer4 marker genes *NECAB1, RORB*; EX42/33/22/32/20, late-born cortical neurons residing in VZ/SVZ/IZ during the first trimester, subsequently migrating to CP by GW15 with elevated expression of the upper-layer marker *CUX2 and CUX1* (**Figs 2B, 2C, S5D, S5E**). In contrast to the increasement of excitatory cells in the SP and CP, the proportion of EX_IN_MIX cells dropped during cortical expansion (**Fig S5A**). Though they exhibited layer preference, only EX_IN_MIX4 located at CP by GW21, most of other cells were still in the IZ, revealing distinct developmental pattern in EX_IN_MIX cells (**Figs 2B, 2C).**

At GW21, the cortical region developed into functional lobes. By calculating gene module via HotSpot^33^, we identified module 17 highly expressed in temporal lobe, and module 18 enriched in the frontal lobe (**Fig 2D; Table S1**). These lobe-enriched gene modules exhibited significant correlations with distinct EX subtypes **(Fig 2E**). Specifically, module 17 showed strong association with EX45, whereas module 18 was predominantly linked to EX11**(Fig 2E**). To investigate the relations between cell type and lobes, we further examined the distribution of cells among different lobes. The excitatory neurons showed lobe specificity, with EX11 is enriched in dorsal lobe, EX33/22/10/45 preferentially distributed in the ventral cortex (**Figs 2F-G, S5F**). To study the function of these region enriched cells, the ligand-receptor interactions between cells in SP and CP were investigated. The NPY signaling pathway involving in food intake, circadian rhythms and memory^34, 35, 36^, abundant in GABAergic neurons, was specifically secreted by EX33 (**Figs 2H, S5G**). The *NPY* peptide and its receptor *NPY1R* both expressed in temporal cortex at GW21 (**Figs 2I, S5H**). The spatial adjacent localization of EX33 secreting *NPY* and EX22/EX2 expressing *NPY1R*, suggested the communication in excitatory neurons though *NPY* signaling pathways (**Fig 2J**). Moreover, EX33 and EX22 exhibited L2/3 neurons identify, while EX2 with L4 neuron feature (**Fig 2C**). Our results indicated that L2/3 EX neurons could modulate not only other L2/3 EX subclusters, but also the L4 EX subcluster (EX2) through *NPY* pathway in the developing temporal cortex.

**Fig.2.**
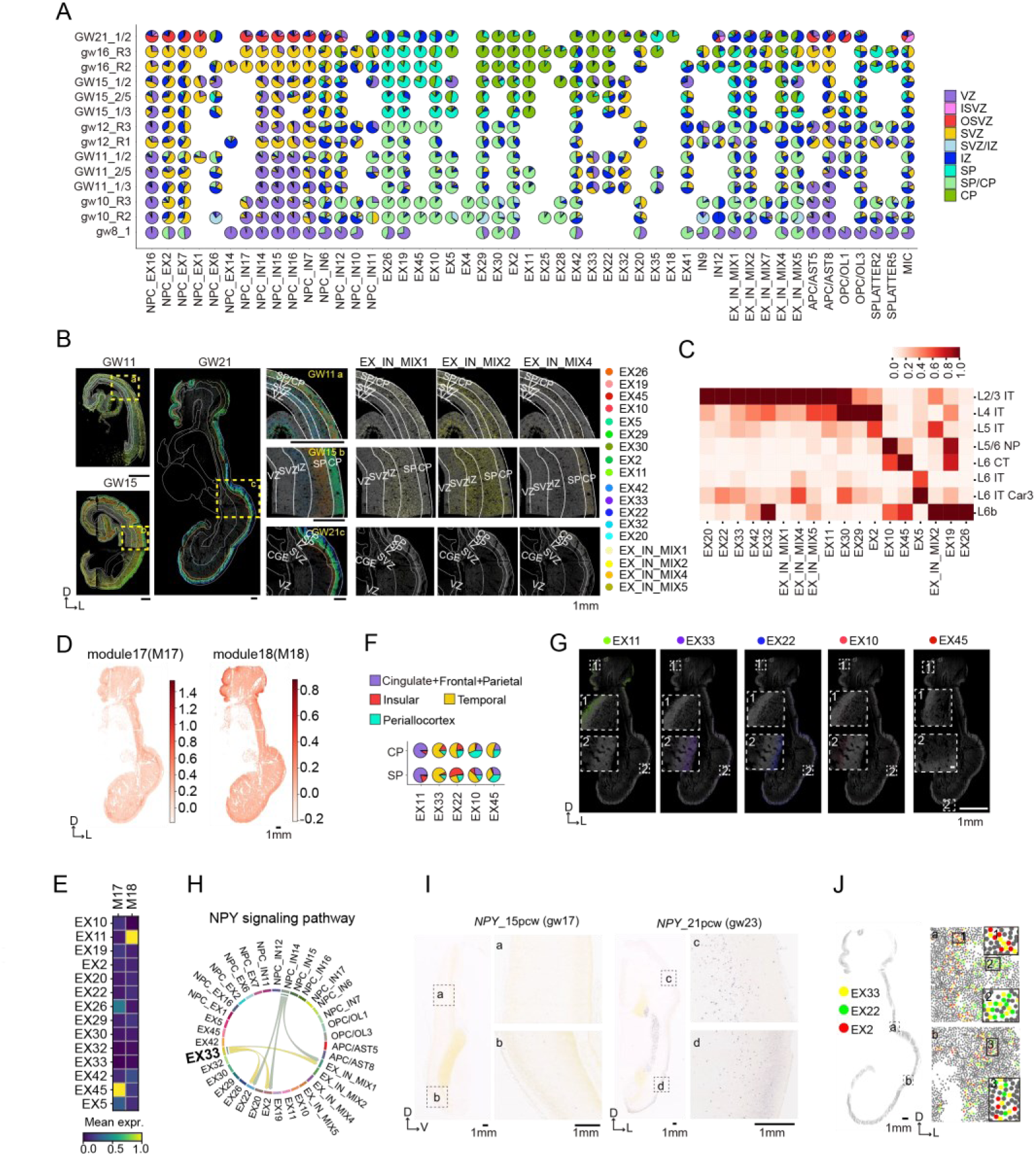
| The dynamic distribution of excitatory cells in cortical expansion and arealization. **A.** Pie chart showing the spatial distribution of cells in cortical subregions during development. GW: samples from this study; gw: published data by Li et al. **B.** Spatial mapping showing the distribution of 14 EX and 4 EX_IN_MIX cell types at single cell resolution at GW11, GW15 and GW21. White solid lines represent the borders of sublayers. D: dorsal; L: lateral. Scales:1mm. C. Heatmap showing the correlation between adult EX and spatially mapped fetal EX. **D.** Spatial visualization of lobe enriched gene modules using Hotspot. Scales: 1mm. **E.** Heatmap showing the correlation of gene modules and EX cell types. F. Pie chart showing the spatial distribution of cells in SP and CP of different lobes at GW21. **G.** Spatial plot showing EX cell types at single cell resolution in different lobes at GW21. White dotted box 1: frontal lobe; White dotted box 2: temporal lobe. D: dorsal; L: lateral. Scales:1mm. **H.** Chord plot showing *NPY* signaling pathway in snRNA using CellChat. **I.** Expression of *NPY* in ISH data at 15pcw and 21pcw from Allen Brain Atlas. Scales: 1mm. D: dorsal, V: ventral, L: lateral. J. Spatial visualization of the expression of ligand *NPY* in EX33 and the receptor *NPY1R* in EX22 and EX2 at GW21 at single cell resolution. Black box represents higher-magnification view. Scales: 1mm. D: dorsal, L: lateral. See also Fig.S5 and TableS1.

### The transcriptomic divergence of the gyrus and neighboring sulcus in the second trimester

To investigate how structural patterning emerges during cortical development, we traced the folding of the neocortex across GW11, GW15, and GW21. The morphological characteristics of sulcal initiation was detected at GW15. At GW11, cortical cell differentiation was uniform along the tangential axis, whereas at GW15 and GW21, maturation trajectories extended from the sulcal to adjacent gyral regions, as revealed by velocity streamlines (**Fig 3A**). For further transcriptomic analysis, the gyri and sulci stereo-seq data were extracted at GW15 and GW21(**Fig S6A, STAR Methods).** A strong gene expression correlation (see **STAR Methods**) was observed between the Sulcus_SP and Gyrus_SVZ (*r* = 0.73), and between the Sulcus_IZ and Gyrus_SVZ (*r* = 0.70) (**Fig 3B**). Regarding the germinal subregions, the Sulcus_SVZ also showed a strong correlation with the SVZ (*r* = 0.77) and IZ regions of the Gyrus (*r* = 0.66); the Sulcus_VZ showed highest correlation with Gyrus_SVZ(*r* = 0.64) rather than the Gyrus_VZ (*r* = 0.53) (**Fig 3B**). These findings demonstrated distinct maturation gradients across cortical compartments: sulcal SP and IZ exhibited delayed maturation relative to their gyral counterparts, whereas sulcal VZ and SVZ mirrored the transcriptional profiles of gyral SVZ.

**Fig. 3.**
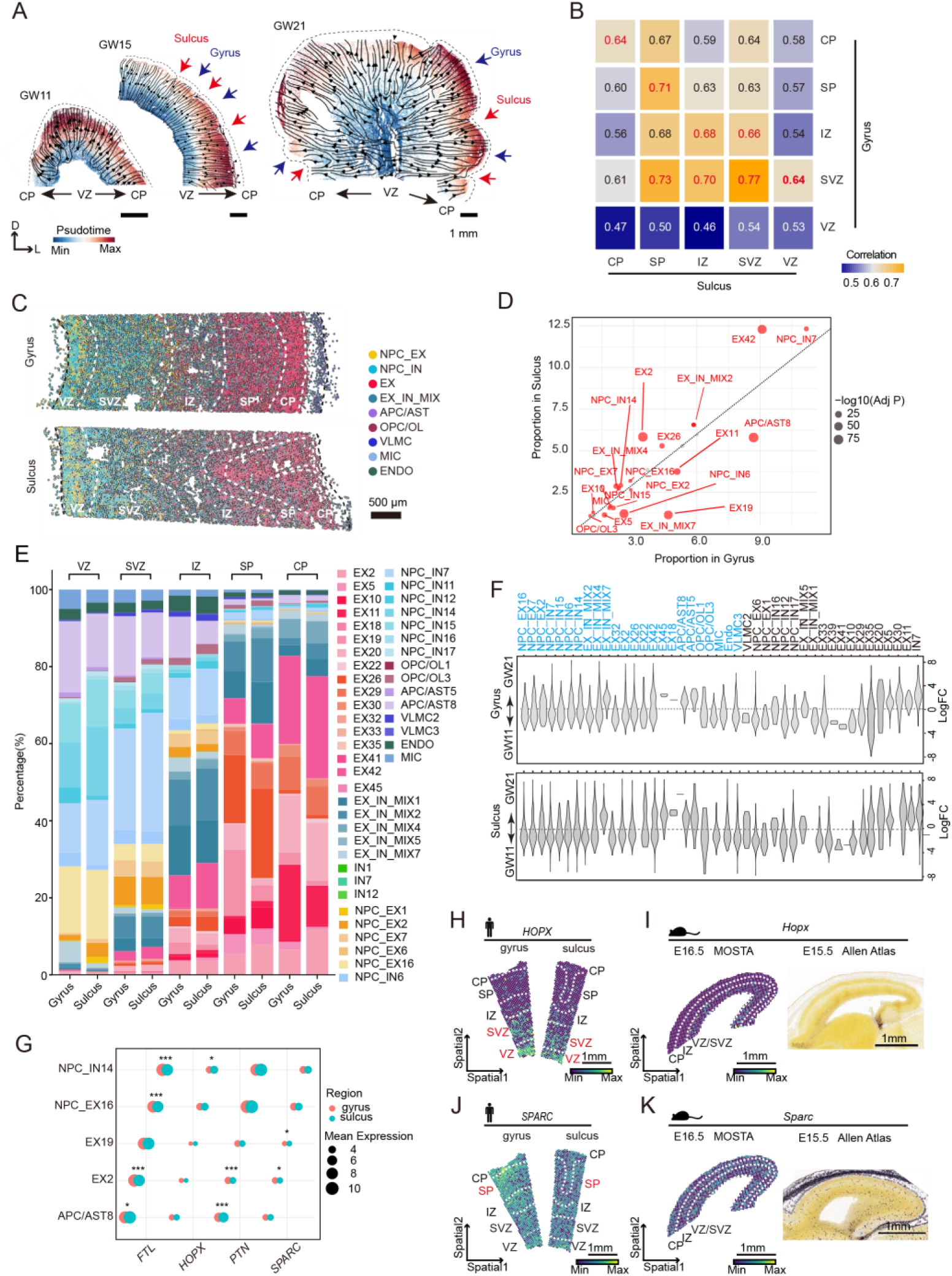
| Spatial heterogeneity of cell types in human cortical folding development. **A.** Pseudotime trajectory analysis at GW11, GW15 and GW21 via SpaTrack. Cortical regions are clustered by FuseMap. Black arrows denote the maturation trend, blue arrows indicate gyri, and red arrows signify sulci. Scales:1mm. **B.** Heatmap showing the gene expression module correspondence between sulcus (x axis) subregions and neighboring gyrus (y axis) subregions. Subregion clusters from FigS6A. **C.** Spatial distribution of 9 major cell types in an example cortex section for gyrus and sulcus at GW15. Scales: 500μm. **C.** Differential cell type proportions between cortical gyri and sulci. Cell types exceeding 1% abundance were shown. Individual points represent cell type-specific proportions. Bubble size corresponds to -log₁₀(adjusted P-value) (Chi-squared test with Benjamini-Hochberg correction). Diagonal line represents equal proportions (y = x). **D.** Stack-bar plots showing the percentage of cell subtypes in each cortical layer between gyrus and sulcus. **E.** MILO analysis showing the proportions of cell types between the gyrus and sulcus across GW11 to GW21. Cell types with statistically significant differential variation patterns (p-value < 0.05, Chi-squared test with Benjamini-Hochberg correction) are highlighted in blue. **F.** Differential expression of *FTL*, *HOPX*, *PTN*, and *SPARC* in selected cell types between gyrus (pink) and sulcus (cyan). Cell types analyzed: NPC_IN14, NPC_EX16, EX19, EX2, APC/AST8. Significance assessed by two-tailed t-test (*p<0.05, **p<0.01, ***p<0.001). **G.** Spatial expression of *HOPX* between gyrus and sulcus at GW15. Scales:1mm. **H.** Spatial expression of *Hopx* in mouse E16.5 cortex sagittal section (*MOSTA data)* (Left). ISH staining of *Hopx* in mouse E15.5 cortex sagittal section (*Allen Atlas*) (Right). Scales:1mm. **I.** Spatial expression of *SPARC* between gyrus and sulcus at GW15. Scales:1mm. **J.** Spatial expression of *Sparc* in mouse E16.5 cortex sagittal section (MOSTA data) (Left). ISH staining of *Sparc* in mouse E15.5 cortex sagittal section (Allen Atlas) (Right). Scales:1mm. See also Fig.S6 and Table S2, S3.

To characterize spatial differences in cell type proportions between the gyrus and sulcus brain regions, we integrated snRNA-seq data with spatial transcriptomic data (**Fig 3C**) and quantitatively analyzed their regional distributions (**Figs 3D, S6B**). Overall, 27 specific cell types proportions in the gyrus different from those in the sulcus, including 4 types of NPC_EX, 5 types of NPC_IN; 7 types of EX, 5 types of EX_IN MIX and 2 types of APC/AST cells (**Fig S6B**). Notably, region-specific enrichment of neuronal subtypes was observed: NPC_EX2, NPC_IN16 (SVZ NPC), EX19 (SP neuron), EX5/EX11 (early-born L4), and APC/AST8 were gyrus-preferring, while NPC_IN7(SVZ NPC), EX26(SP neuron), EX2 (early-born L4), EX42 (late-born L2/3), and EX_IN_MIX2 were sulcus-enriched (**Figs 3D, S6B**). Subregional analysis revealed gyrus-to-sulcus gradients in cell-type proportions across all cortical layers (VZ, SVZ, IZ, SP, CP; **Fig 3E, Table S2**). APC/AST8 showed gyrus-biased distribution spanning VZ to IZ, while NPC_IN7 exhibited peak enrichment in SVZ and EX42 demonstrated maximal CP divergence with sulcus bias. Temporal quantification across developmental stages (GW11–GW21) revealed dynamic, region-specific shifts in NPCs (NPC_EX2,7,16; NPC_IN6,7,14,15), excitatory neurons (EX2,18,22,26,32,42,45), and glial/non-neuronal cells (APC/AST5,8; OPC/OL1,3; MIC; VLMC) (**Fig 3F**).

To understand the contribution of regulatory factors to the asynchronous development of the sulci and gyri, the profile of differential gene expression in various sublayers between gyrus and sulcus were compared (**Figs S6C, D; Table S3**). A total of 48 DEGs were identified in the VZ, 50 in the SVZ, 50 in the IZ, 128 in the SP, and 194 in the CP (**Table S3**). Among the upregulated DEGs in gyri, several genes were associated with neurogenesis and synapse organization, such as *HOPX, PTN, VIM, and TUBB2A*^37, 38, 39, 40^, while others were linked to the functions of oligodendrocytes/astrocytes, including *SPARC* and *GFAP*^41, 42^ (**Fig S6C, S6D**). To elucidate the patterns of cell type-specific expression in the DEGs, we analyzed the profile of gene expression across distinct cortical cell types by using stereo-seq datasets from the gyrus and sulcus (**Fig 3G**). The analysis revealed that *HOPX* and *SPARC* were predominantly expressed in APC/AST and NPC_EX (including radial glia cells) (**Fig S6E**). However, *HOPX* exhibited significantly higher enrichment in gyrus-localized NPC_IN14 rather than any NPC_EX subclusters compared to its sulcus counterpart. *PTN* showed elevated expression in both NPC_IN14 and NPC_EX16, with gyrus-localized EX2 and APC/AST8 displaying higher *PTN* expression than their sulcus counterparts. Furthermore, *FTL* was highly expressed in NPC_IN4, NPC_EX16, EX19, EX2, and APC/AST8 within the gyrus (**Fig 3G**). These findings highlight region-specific molecular signatures across neuronal and glial cell populations during cortical development.

The spatial distribution of DEGs related to cortical folding was analyzed in human fetal brains and smooth-brained species (**Figs 3H-K**). The gene expression patterns in the developing human fetal brain at GW15 were compared with those at equivalent developmental stages (E15.5-E16.5) of the mouse cortex^43, 44^. In human GW15 cortex, *HOPX* was highly expressed in the VZ/SVZ, particularly in gyral VZ (**Fig 3H**), while mouse *Hopx* (E15.5-E16.5) showed uniform, sparse VZ distribution, in the Atlas of the Developing Mouse Brain (*MOSTA*) data^45^ (**Fig 3I**). *SPARC* displayed broad human cortical expression but higher gyrus-specific subplate enrichment (**Fig 3J**), contrasting with its diffuse A-P gradient in mice (**Fig 3K**). Similarly, *ID2*, *ZEB1*, and *VIM*—human gyrus-sulcus markers— exhibited A-P gradients in E14.5/E16.5 mice (**Figs S6E-F**).The distinct but evolutionarily conserved spatial expression patterns of these genes in human and mouse cortices suggest their potential involvement in the development of cortical folding structures.

### The characteristics of D1/D2 hybrid MSNs in the developing striatum

Medium spiny neurons (MSNs) were isolated from interneurons through sample library identifier-based segregation using datasets from both Shi et al.^15^ and our independent experiments. snRNA-seq analysis of 18,024 cells revealed five transcriptionally distinct clusters classified as MSNs (**Figs 4A, S7A-B**). Notably, two clusters demonstrated concurrent expression of D1- and D2-type dopamine receptor markers, identifying a hybrid D1/D2 MSN subpopulation. Transcriptomic co-expression of prototypic markers *TAC1* (D1-associated) and *PENK* (D2-associated) was validated across both snRNA-seq and ST datasets, further corroborating the existence of this hybrid MSN population **(Figs 4B, S4C**). Spatial transcriptomic analysis demonstrated significant spatial co-expression of *TAC1* and *PENK* transcripts across striatal subdomains, indicative of a pronounced enrichment of D1-D2 co-expressing MSNs within striatal compartments relative to GE-derived neuronal populations (**Figs 4B, S4D**). Regional specificity was evident, with *TAC1* exhibiting a marked enrichment in ventromedial striatal territories, whereas *PENK* expression was predominantly localized to dorsomedial striatal regions. Over time, the prevalence of D2-MSNs exhibits a progressive increase, whereas the proportion of D1/D2 hybrid populations demonstrates a concomitant decrease (**Fig S4D**). This dorsoventral stratification suggests divergent expression gradients of D1 and D2 MSN subtypes, further supporting functional and molecular heterogeneity within striatal circuits. These results provide direct evidence for the existence of a D1/D2 hybrid MSN subpopulation and highlight its anatomical predominance in the striatum. snRNA-seq analysis revealed distinct spatial enrichment patterns of D1/D2 hybrid MSN subtypes: D1/D2 hybrid MSN1 exhibited pronounced localization within the GE, while D1/D2 hybrid MSN2 demonstrated preferential enrichment in the striatum. During gestational maturation, the proportional abundance of D1/D2 hybrid MSN1 subtypes exhibited a progressive decline within both the GE and striatum, whereas the relative frequency of D1/D2 hybrid MSN2 subtypes showed a concomitant increase across both regions with advancing gestational age **(Figs 4C, S4E-F)**. These region-specific distributions were further validated through integrative spatial mapping of snRNA-seq clusters onto ST datasets, which corroborated the anatomical segregation of D1/D2 hybrid MSN1 and D1/D2 hybrid MSN2 subtypes across the GE and striatum, respectively **(Figs 4D and 4E)**.

**Fig. 4.**
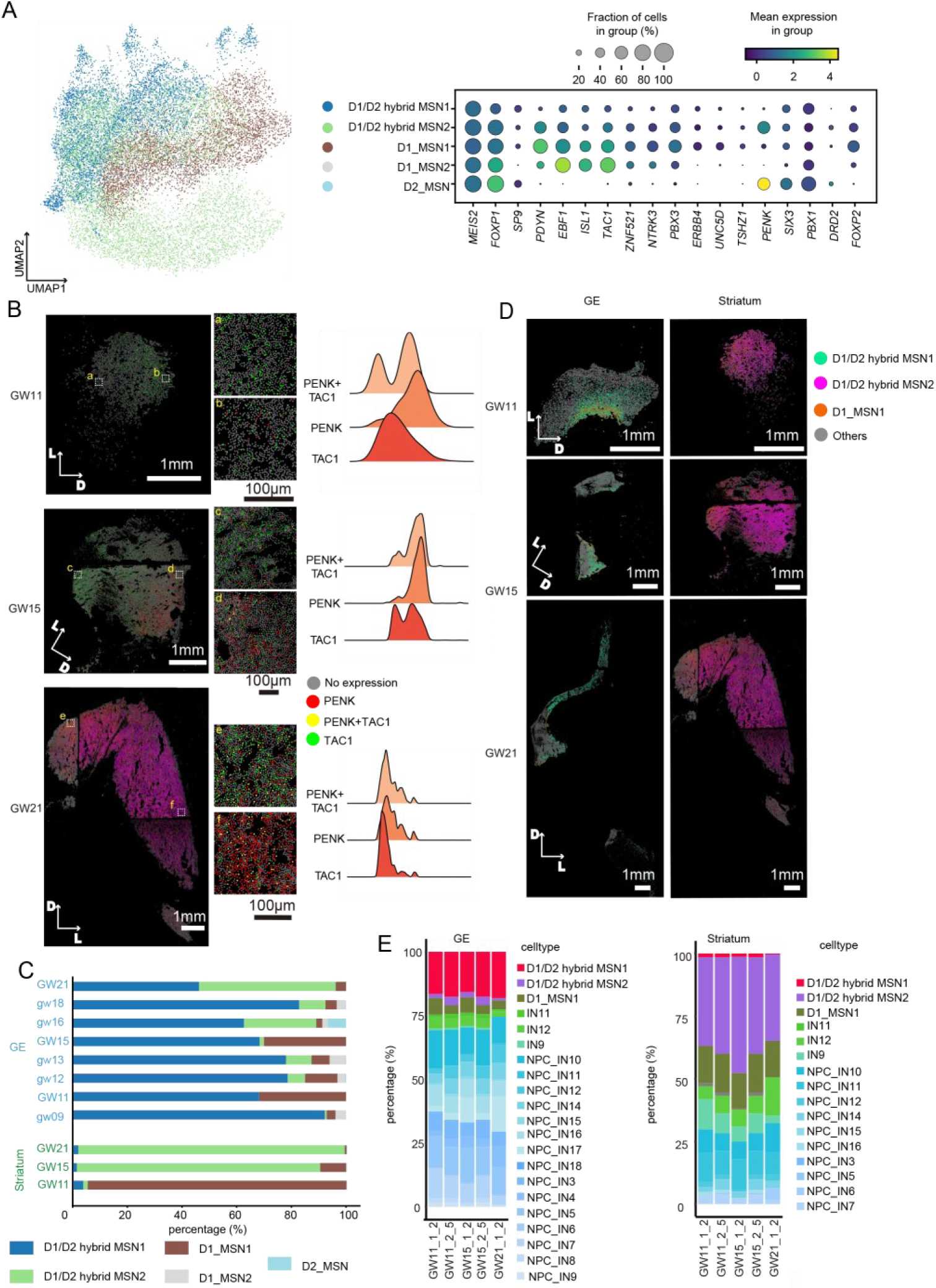
| Spatiotemporal dynamics of D1/D2 hybrid MSNs. **A.** UMAP and dot plot showing clustering and specific markers of MSNs. Left: UMAP projection of MSNs isolated from integrated snRNA-seq data (see Fig. 1B), partitioned into five transcriptionally distinct clusters. Right: Dot plot depicting cluster-specific marker expression. Two clusters were classified as D1/D2 hybrid MSNs (co-expressing canonical markers), two as D1_MSNs, and one as D2_MSNs. Bubble size shows the percentage of cell expressed; scale bar shows the average expression level. **B.** Spatial co-expression showing neuropeptide markers expression in striatal regions. Left: Representative spatial expression patterns of *TAC1* and *PENK*. Right: Ridge plot quantifying *TAC1*, *PENK*, and co-expression signal density along the ventral-dorsal axis. D: dorsal, L: lateral. Scales: white = 1 mm, black = 100 μm; inset shows higher-magnification view. **C.** Bar plot showing the proportional distribution of the five MSN clusters within the GE and striatum at distinct gestational weeks using snRNA data. GW: samples from this study; gw: published data by Shi et al. **D.** Integration of snRNA-seq and spatial transcriptomics data showing cell type distribution patterns in GE and STR. D: dorsal, L: lateral. Scales: 1mm. **E.** Stack-bar plot summarizing the regional and gestational stage-specific distribution of spatial cell types across ST sections in GE and striatum. See also Fig. S7.

### The formation of patch and matrix compartments in the striatum

The striatum is heterogeneously consisted of patch and matrix compartments, which are distinguished by their distinct neurochemicals^46^. To explore the spatial expression pattern in the developing striatum, we performed polynomial regression on the normalized gene expression data at each time point, resulting in 46 upregulated and 46 downregulated genes (**Fig S8A**). *PCP4, PCDH17* and *PENK showed* higher expression on dorsal-lateral side (**Fig S8B**). To explore the dorsal-lateral versus ventral-medial genes of striatum, spatial variable gene (SVG) analysis was performed on the LGE-mantle zone (promising striatum) on coronal sections. Transcriptional profiling revealed distinct spatial enrichment patterns, with 4 genes exhibiting significant enrichment in the ventromedial region, while 25 genes showed pronounced enrichment in the dorsolateral region, suggesting region-specific functional specialization (**Fig S8C)**. Some genes (eg. *NRGN, NTM)* maintained region preference across all time points, whereas other genes (*GUCY1A1, CRABP1* and *SIX3*) did not (**Fig S8C**). At GW21, patch and matrix-like compartments were observed (**Figs 5A and S8D**). Notably, *GABRA5* and *EPHA4*, the known patch and matrix markers in the adult human brain^47^, colocalized with patch-enriched and matrix-enriched SVGs, respectively (**Fig S8E**). The patch and matrix marker genes exhibited compensatory expression, highlighting the unique features of these compartments in the striatum.

**Fig. 5.**
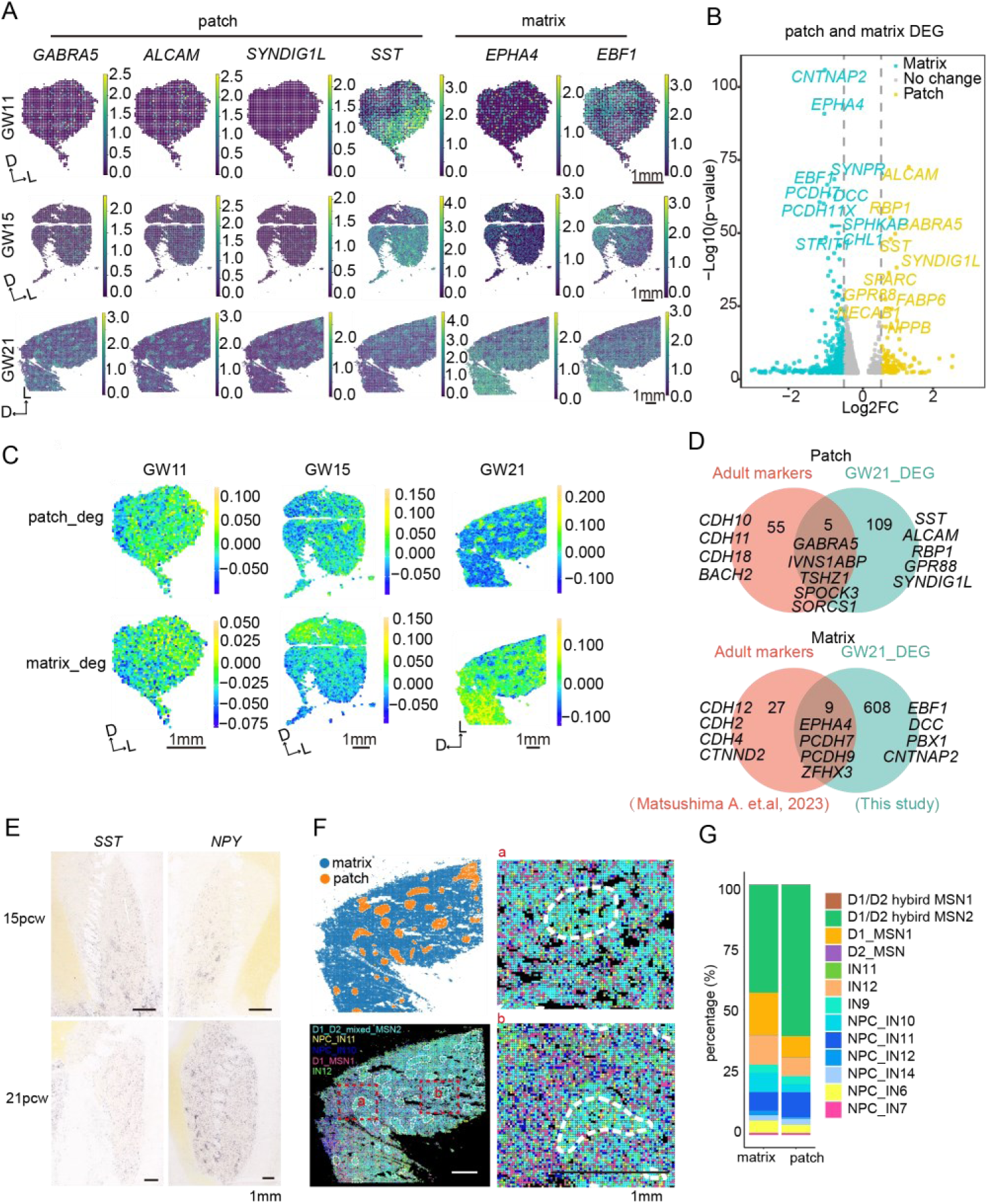
| Development of striatal patch and matrix compartments. **A.** Spatial visualization of the spatial variable genes showing the structure of striatal patch and matrix at GW11, GW15, and GW21. D: dorsal; L: lateral. Scales: 1mm. **B.** Volcano plot showing DEGs of patch and matrix at GW21. **C.** Spatial visualization of patch and matrix gene module expression at GW11, GW15 and GW21. D: dorsal; L: lateral. Scales: 1mm. **D.** Venn plot showing the DEGs of patch and matrix shared by this GW21 spatial data and published adult human striatum data. **E.** The expression of *SST* and *NPY* in ISH fetal striatum at 15pcw and 21pcw from Allen Atlas. Scales:1mm. **F.** Spatial-ID showing spatially resolved single cell types of patch and matrix. Identified region of patch and matrix by Gaussian blur (upper left). Spatial visualization of enriched cells types in patch and matrix (lower left). Magnification of mapped cells (right). White dotted lines are the borders of patch and matrix. Red dotted lines are zoom-in zones. D: dorsal; L: lateral. Scales: 1mm. **G.** Stack-bar plot showing the percentage of cell types in patch and matrix. **See also** Figs. S8, S9, S10 **and Table S4.**

To further analyze the developmental features of the patch and matrix compartments, we applied Gaussian blur to extract patch and matrix (**Fig S8F;** see **STAR Methods**), resulting in 114 patch markers and 617 matrix marker genes (**Fig 5B; Table S4**). Pathways related to the response to thyroid hormone stimulus, synapse organization, innervation and regionalization were enriched in the patch. Conversely, in the matrix, the enriched GO terms were primarily related to adhesion molecules, axon regeneration and neuronal projection, indicating cell migration and axon development (**Table S4**). These results suggested that distinct biological processes occur in the patch and matrix compartments. Gene module scores for both patch and matrix spatial gene sets based on DEGs were calculated across all time points (**Fig 5C**). The patch module demonstrated ubiquitous distribution throughout the dorsolateral striatum by GW 11 and progressively organized into distinct patch compartments at GW15 and GW21. In contrast, the matrix module exhibited a complementary spatial distribution pattern to the patch module by GW21, consistent with established striatal compartmentalization principles (**Fig 5C**).

To investigate developmental-specific features as well as conserved genes in patch and matrix compartments, we compared patch and matrix markers with published data from the adult brain^47^ (**Fig 5D**). Remarkably, *CDH* family was enriched in the adult patch and matrix but not in the fetal brain. Only 5 patch genes and 9 matrix genes were consistently related to the maintenance of the patch or matrix compartments. A total of 109 genes, such as *SST*, were enriched in developmental patches (**Fig 5D**). The temporal features of top 40 developmental patch genes were plotted (**Fig S9A**). Importantly, at GW15, *SST* exhibited a patch-like pattern, while other genes, such as *SYNDIG1L,* did not (**Figs 5A, S8D, S9A**). These findings suggest that the genes in patches gradually and asynchronously developed. Furthermore, *SST* was expressed in 98.1% of the identified patch cells at GW11 and 100% at GW15. Although the percentage decreased to 71.4% at GW21, it still remained above the average level of all genes (**Figs S10A and S10B**). Moreover, the expression patterns of *CORT* and *CENPS-CORT (*both encoding cortistatin structurally similar to somatostatin), as well as *NPY,* were all similar to *SST* at GW15 and GW21 (**Fig S10C**). The expression of *SST* and *NPY* exhibited a patch-like pattern in ISH data of fetal striatum at 15pcw and 21pcw (**Fig 5E**). These findings indicated that *SST* is a patch-enriched gene in the human fetal brain, and these patch and matrix region-specific genes dynamically contribute to and maintain the properties of the developing striatum. We then explored patch and matrix genes in mouse embryo **(Fig S10D)**. *Gabra5* and *Gpr88* exhibited patched pattern at E18.5, *Alcam* at P4, however, *Sst* and *Npy* did not show patch enriched characteristics (**Fig S10D**). These suggested that despite the structures are conserved, the genes in the developing patches are different between human and mice.

Subsequently, the spatial distribution of single cell types in patch and matrix were quantified (**Figs 5F,5G**). At GW21, the proportion of D1/D2 hybrid MSN2 and NPC_IN11 were higher in patch, while D1_MSN1, NPC_IN10 and IN12 were greater in matrix (**Figs 5F, 5G**). Though at low levels, *SST/NPY/CORT* were co-expressed in the NPC_IN11 cells (**Fig S10E**). These results suggested neuropeptide *SST* and *NPY* may involve in patch formation. Overall, our results demonstrated that the patch and matrix undergo dynamic development.

## Discussion

This study established a single-cell resolution spatial map of developing human brain from GW8 to GW21. By systematically analyzing the spatiotemporal gene expression and cellular distribution patterns, we reveal new mechanisms governing cortical expansion and uncover previously unrecognized developmental principles orchestrating striatal organization within the subpallium.

### Cortical expansion and arealization

The developing cortex is anatomically categorized as the VZ, SVZ, IZ, SP and CP. SP was observed as a distinguished layer at GW10 after the emergence of the CP, which occurred at approximately GW8-GW9^3^. In addition to the concept of SP based on immunostaining, this study identified the subplate and its cellular composition at the transcriptional level **(Figs 2A, S5A**). The SP was separated from the preplate SP/CP after GW12. SP neurons in the SP layer are the earliest generated and mature neurons in the cerebral cortex, guiding neuronal migration towards the CP and establishing the earliest synapses^48, 49^. Several studies have characterized human transcriptional features of SP-specific neurons in the human fetal brain^20, 50, 51^.The neurons (EX26/19/45/10 and EX_IN_MIX2) largely constituted the SP exhibited strong correlation with adult L5/6 neurons, align with findings that SP neurons differentiate into deep layer cortical neurons^52^, suggested a correlation of SP with layer 6 in adult brain. The cells in the subplate are dynamically changed during cortical expansion as more interneurons migrating to the cortical plate^32^. To elucidate the development of subplate and spatiotemporal segregation mechanisms between the subplate and layer 6, systematic analysis of fetal cortical samples across sequential developmental stages is required. Coinciding with the clear anatomical separation of SP and CP, deep and upper layer neurons underwent progressive laminar segregation concurrent with the emergence of distinct cortical lobes at GW21 (**Figs 1C, 2B**). Previous studies have identified that *NPY* is predominantly expressed in excitatory lineage cells within the parietal, temporal, and V1 cortices compared to the PFC, motor cortex, and somatosensory cortex^53^. However, these studies did not specify the neuronal subtypes expressing *NPY* and their laminar distribution, or their potential connectivity with other neuronal populations. Here, we identified a temporal lobe-enriched EX subtype, EX33, characterized by enriched *NPY* expression, residing in layer 2/3. This subtype exhibits potential functional interactions with L4 neuron EX2 through *NPY-NPY1R* ligand-receptor pairs (**Figs 2F-I, S5G**). The *NPY* plays critical roles in neural systems, such as promoting proliferation of postnatal neuronal precursor cells^54^, regulating anxiety and stress related responses^55^, feeding^56^, learning and memory^57, 58^, endocrine function^59^, and circadian rhythms^60^. The function of *NPY* in L2/3 EX neurons remain to be addressed. In adult macaque V1 cortex, a type of EX neuron expressing NPY was also found restricted to L2/3^61^. Yet, the role of the *NPY*-expressing EX in cortical arealization remain understudied. In the developing mouse cortex, *Npy* expression becomes detectable post-E18.5, with higher expression levels observed in anterior cortical regions compared to posterior regions until postnatal day 4 (P4). By P14, *Npy* expression becomes uniformly distributed along the anterior-posterior axis, persisting through P28. (https://developingmouse.brain-map.org/). The spatiotemporal expression pattern of *Npy* in the developing murine cortex is divergent from that observed in the human fetal cortex. These findings highlight the need for stratified analyses of *NPY*’s functional roles across species and cortical layers.

### Transcriptomic features of developing fissures

Recent studies have shown that specific molecules or modeled mechanicals promote the cortical folding in gyrencephalic species^62, 63, 64, 65^. This study provided transcriptome resources for the patterns of gene expression in various developing fissures using the human fetal brain stereo-chip (**Figs 3A, S6A**). It was the first systematic identification of DEGs and specific celltypes from the gyrus and its adjacent sulcus in the human fetal cortex in situ.

In human cortical folding, multiple sulci (e.g., Superior Frontal Sulcus, Callosal sulcus, Cingulate Sulcus, Sylvian fissure) and adjacent gyri were included for analysis (**Fig S6A**). While gene expression differences between gyri and sulci were subtle (**Fig S6C**), key divergences were observed in NPC/APC/EX ratios—particularly in SVZ and SP regions (**Figs 3D, 3E, Table S2**). The gyrus exhibited significantly more *HOPX*-enriched APC/AST8 cells (**Fig 3G**), likely due to: (1) higher APC/AST8 and NPC_IN14 proportions (major factor), and (2) elevated *HOPX* expression in gyral NPC_IN14 (secondary factor).

### The characteristics of D1/D2 hybrid MSNs in the developing striatum

D1/D2 hybrid MSNs have been documented in multiple species, including primates (e.g., macaques), rodents, and other model systems, across numerous studies^66, 67, 68, 69, 70^ However, their presence during human neurodevelopment remains unconfirmed. In this study, we provide the first evidence of D1/D2 hybrid MSNs in the developing human brain, by identifying these cells across distinct developmental stages with striatal enrichment observed from GW15 onward. We further classified two subtypes of D1_D2 co-expressing MSNs: D1/D2 hybrid MSN1 exhibited preferential localization to the GE, whereas D1/D2 hybrid MSN2 demonstrated pronounced enrichment in the striatum. While a comparative analysis of these distinct clusters falls outside the scope of this investigation, their phenotypic and functional characterization represents a critical avenue for future research.

### Transcriptomic and cellular dynamics in the formation of patch and matrix compartments

A previous study has demonstrated that striatal neurons in patch and matrix compartments undergo distinct developmental processes^9^. Comparative analysis with prior studies of the adult human striatum^47^ revealed evolutionarily conserved genes enriched in striatal patch (e.g., *GABRA5*) and matrix (e.g., *EPHA4*) compartments. Additionally, we identified novel developmental-stage-specific markers (*ALCAM, GPR88,* and *SST*), which to our knowledge represent the first report of their dynamic expression patterns during striatal ontogeny (**Fig 5D**). Notably, *SST* was well colocalized within all patch compartments and was enriched in patches as early as GW11 (**Fig 5A**). Spatial profile of striatal cell subtypes showed compartment-specific enrichment, with NPC_IN11 interneurons (co-expressing *SST*, *NPY*, and *CORT*) exhibiting a significantly higher prevalence within the patch compartment compared to the matrix. Moreover, the *NPY* showed a patch like pattern at GW15 and pcw21 from ISH data. However, the *SST* and *NPY* didn’t show a patch enriched pattern in developing mouse brain, and there was no difference in the density of *SST* between the patch and matrix in the adult mouse striatum^71^. These data implicate *SST* and *NPY* in human regulatory mechanisms underlying striatal compartmentalization, contrasting with murine models where these neuropeptides lack analogous roles. The spatiotemporal dynamics of patch-matrix developmental remodeling and the contributions of *SST/NPY* signaling to this process remain poorly characterized, necessitating systematic interrogation across postnatal-to-adult transitions. In summary, our study depicted a spatial-temporal map of developing regions in the brain by stereo-seq and snRNA-seq. We demonstrated dynamic structural changes associated with cortical layer expansion, gyrification, and subpallial striatal compartment development. This study provides novel insights into the development of the cortex and striatum and enhance our overall understanding of brain development.

### Limitations of the study

The efficient capture of all genes during the experimental process was challenged by using Stereo-seq, particularly for those genes with low expression levels. This may lead to the inadvertent exclusion of significant genes from both processes of experimental and data analysis. Cells in close proximity may be erroneously binned together to increase statistical power of analysis. Due to the scarcity of human fetal brain samples, the obtainment of multiple replicates was not feasible for this study. Further experimentation is necessary to test and confirm the conclusions and hypotheses generated from this study.

## Methods

### HUMAN SUBJECTS

Human embryo tissues were obtained in accordance from West China 2^nd^ University Hospital with ethic approval of Ethics Committees of West China 2^nd^ University Hospital (19H1194). All women gave written consent to the use of fetal tissues according to the ISSCR guidelines for fetal tissue donation. Fetal brain samples were collected after the donor patients signed an informed consent document that was in strict observance of the legal and institutional ethical regulations for elective pregnancy termination specimens at West China 2^nd^ University Hospital. The gestational age was measured in weeks from the first day of the woman’s last menstrual period.

### Tissue collection

Fetal brains were collected in ice-cold artificial cerebrospinal fluid (ACSF) containing 125.0 mM NaCl, 26.0 mM NaHCO_3_, 2.5 mM KCl, 2.0 mM CaCl_2_, 1.0 mM MgCl_2_, 1.25 mM NaH_2_PO_4_at a pH of 7.4 when oxygenated (95% O_2_ and 5% CO_2_). The brain hemispheres were collected from three fetuses (11 gestational weeks, GW11; 15 gestational weeks, GW15; 16 gestational weeks, GW16; 21 gestational weeks, GW21). The isolated brain hemispheres were quickly wiped dry with sterile gauze and mixed thoroughly with 4°C optimal cutting temperature (OCT) compound (4583#, Sakura). Subsequently, the brain blocks were transferred to a metal mold fully embedded with 4°C OCT, quickly frozen on dry ice, and then stored in -80°C refrigerator. To minimize RNA degradation, all solutions were prepared with sterilized water containing diethyl pyrocarbonate (DEPC), and all instruments were washed with sterilized water containing DEPC and RNase Zap (AM9780, Invitrogen). The whole tissue collection process was completed within 30 minutes.

### Sample collection

The 1/3, 2/5 and 1/2 coronal sections of GW11 and GW15 fetal brain, and 1/2 coronal section of GW21 fetal brain were collected for stereo-seq analysis. We aligned 4 chips to form a square and the 2/5 coronal section of GW15 was attached on the top. The 1/2 coronal section of GW15 was attached on the top of 3 aligned chips, and the 1/2 coronal section of GW21 was attached on the top of 14 aligned chips (**Figs 1C, S2C**). Each chip was then used for stereo-seq analysis. The adjacent tissue of each coronal section, except 1/3 section of GW15, was collected for snRNA analysis. In addition, the following tissues were also collected for snRNA analysis: the cortex of 1/3 coronal section of GW11, 1/3 coronal section of GW16, 1/2 coronal section of GW15 as well as four functional groups of cortex of 1/2 coronal section of GW21; the GE region on 1/2 coronal section of GW11 and GW15, and LGE and MGE region on 1/2 coronal section of GW21; the striatum on 1/2 coronal section of GW11, GW15 and GW21 respectively.

### Tissue-section preparation for snRNA-seq

The DNBelab C Series Single-Cell Library Prep Set (MGI, 1000021082) was used as previously described^72^. Briefly, single-nucleus suspensions were used for droplet generation, emulsion breakage, bead collection, reverse transcription and cDNA amplification to generate barcoded libraries. Indexed libraries were constructed according to the manufacturer’s protocol. The sequencing libraries were quantified by Qubit ssDNA Assay Kit (Thermo Fisher Scientific). The sequencing libraries were sequenced by the DIPSEQ T7 sequencer at BGI-Shenzhen with the following sequencing strategy: 41-bp read length for read 1, and 100-bp read length for read 2.

### Tissue-section preprocessing for Stereo-seq

Stereo-seq library preparation and sequencing were performed as previously described^73^. Briefly, tissues were adhered to the Stereo-seq chip. A nucleic acid dye staining (Thermo Fisher, Q10212) was performed to assess tissue morphology and quality. Tissue sections were permeabilized at 37° for 12 min and reverse transcription was conducted by second strand synthesis and cDNA denaturation. The cDNA was purified using AMPure XP beads (Vazyme, N411-03). The indexed single-cell RNA-seq libraries were constructed according to the manufacturer’s protocol. The sequencing libraries were then sequenced by the DIPSEQ T7 sequencer at BGI-Shenzhen with the following sequencing strategy: 50-bp read length for read 1, and 100-bp read length for read 2.

### snRNA data quality control and pre-processing

The sequencing data were processed as previously described^11^. Nuclei with Unique Molecular Identifier counts (UMIs) counts <1500, or gene counts <500, or mitochondrial gene count percentage >20% were removed. Ribosomal genes, mitochondrial genes and blood-related genes (*HBA1, HBA2, HBB, HBG1, HBG2, HBQ1, HBD, HBM, HBE1 and HBZ*) were also removed according to published paper^74^. Data were log-normalized and top 3000 highly variable genes were selected using Seurat^75^ LOESS method for downstream analysis.

### snRNA data integration for dimension reduction, clustering and subclustering

The single-nuclei data from different sources^12, 14, 15^ were integrated with data from this study, and the conservative gene sets of each single-nuclei data set were used for downstream analysis. The batch effects were corrected using harmony algorithm, and the data dimension was reduced and projected to UMAP. Cells were divided into three major categories based on the marker genes of the main excitatory neurons, inhibitory neurons and non-neuronal neurons. Subclustering was performed on each major cell group. Wilcoxon rank sum test was applied using the “sc.tl.rangk_genes_groups”function for differential gene expression analysis among clusters. Genes with log2 fold-change > 1 and FDR < 0.05 were retained as significant marker genes for the subcluster. Jaccard similarity was calculated with the top 50 ranked marker genes of each pair of cell clusters, and clusters were combined with Jaccard similarity larger than 0.8. 0.1% of the cells were identified as outliers and removed based on the cell distance to the center point. Subclustering analysis identified 136 distinct subtypes, which were subsequently aggregated into 12 biologically defined cell types based on their marker gene expression profiles.

### MILO analysis

For snRNA, after data integration, the cells were then clustered to three trimesters and the differential abundant analysis was performed using python MILO package (v0.1.1). For ST data, after cell mapping, the mapped cell types in ST data were quantified and the same analysis was performed.

### Cross-method correlation analysis

Gene specificity correlation analysis was used to evaluate the concordance between SpaGCN and FuseMap annotations. A unified set of 2,000 highly variable genes (HVGs) was identified from the integrated SpaGCN-FuseMap dataset using FindVariableFeatures function (Seurat; selection method = ‘vst’, mean-variance threshold = 1.5). Gene specificity vectors between homologous subregions (e.g., VZ_FuseMap vs. VZ_SpaGCN) were compared using rank-based Spearman correlation. Statistical significance was determined through two-tailed permutation testing (10,000 iterations) with Benjamini-Hochberg correction for multiple hypothesis testing.

### Gyrus-Sulcus subregions Pairwise Cluster Correlations

The stereo-seq data of gyrus/sulcus regions were lassoed by their anatomically spatial structure, including Superior frontal sulcus (SFS), Callosal sulcus (CaS), Cingulate sulcus (CiS), Sylvian fissure (SF)^76, 77^, provisional prigeminal fissure (PPF)^78^, and their adjacent gyri. For gyrus/sulcus profiling, we manually lassoed three paired gyral and sulcal regions from anatomically defined 1/3 anterior, 2/5 middle, and 1/2 posterior coronal sections of GW15 brains. Two gyral and three sulcal regions were segmented from GW21 mid-posterior coronal sections, with technical replicates (n=4 or 5 sections per region) (Figs S6A). The region of interest was selected on ssDNA (using Adobe Photoshop v25.3.1). Then the expression matrix was aligned to ssDNA and the region of interest was lassoed. The subregions (VZ/SVZ/IZ/SP/CP) were annotated by FuseMap. The subregion gene expression was normalized using the SCTransform function in Seurat. The correlation analysis between gyrus and sulcus consisted of:1) the selection of a common gene set for the comparison. Variable genes identified from the gyrus and sulcus combined data using Seurat FindVariableFeatures function (selection.method = ‘vst’, nfeatures = 2,000) were used to generated averaged scaled expression values and correlated. 2) the calculation of Spearman rank correlations (and their p-values) of gene specificities among all pairs of subregions in the gyrus and sulcus.

### Comparative analysis with published atlas

Reference subregional definitions^12^ were reprocessed using FuseMap-based harmonization.

Transcriptomically matched subregions were identified through reciprocal nearest neighbor alignment (integration weight = 0.7) in PCA-UMAP latent space (dims=1:30). Subsequent correlation analyses followed the aforementioned Spearman framework.

### Transfer fetal brain celltype to adult brain celltype

XGBoost (eXtreme Gradient Boosting) model^79^ was trained to transfer our fetal brain cell type annotation to adult brain single-cell atlas. The model objective was set as multi-classification task (evaluation metrics: mlogloss, learning rate=0.001). The model was trained on 1 Nvidia A100-80G GPU.

### Sanky plot

The ST data at same gestational week were combined, and the data from adjacent gestational weeks were used to calculate Jaccard similarity and z-score. R package networkD3 (0.4) was then used to generate Sanky Plot.

### Differential expression genes (DEGs) and GO enrichment analysis

Seurat FindAllMarkers function was used to calculate DEGs with minimum expression percentage as 0.25. The other parameters were set as default. Calculated DEGs with adjusted p values > 0.5 were filtered out, and the remaining DEGs were ordered by descending average log_2_FC value. ClusterProfiler^80^ package and org.Hs.eg.db^81^ package were used to calculate enriched GO categories based on the NIH database tool DAVID (v6.7) (https://david.ncifcrf.gov)^82, 83^.

### Ligand-receptor communication analysis

CellChat^84^ package was used to calculate the ligand-receptor communication of mapped cells.

### Spatial data (ST data) pre-processing

Expression profile of matrix of human brain was divided into non-overlapping bins covering an area of 100*100 DNB to form bin100 data (∼50 µm) and the transcripts of the same gene were aggregated within each bin. For GW15 and GW21 chips, the chips were aligned according to chip coordinates respectively.

### Cluster ST data using SpaGCN

Spatial data of bin100 were clustered using SpaGCN^17^ for each coronal section. The resolution and region identification were set according to Brainspan atlas of the developing human brain (https://www.brainspan.org/static/atlas).

SpaGCN was employed to identify spatial domains jointly by using a graph convolutional neural network to learn a representation aggregating gene expression data from surrounding spots to identify the domains. The adjacency graph used in the convolution was constructed based on spatial location. The resolution and region identification were set according to Brainspan atlas of the developing human brain (https://www.brainspan.org/static/atlas).

### Lasso the region of interest in ST data

The region of interest was selected on ssDNA. Then the expression matrix was aligned to ssDNA and the region of interest was lassoed.

### Spatial Transcriptomics cell segmentation

The optimal cell segmentation model (cell_segmentation_v3.0.onnx) developed by Stereopy^85^ was applied for Stereo-seq data with extremely dense cell populations to perform nuclear segmentation on human fetal brain nucleic acid staining images. Subsequently, the segmented cell masks were integrated with Stereo-seq data using Stereopy’s stereo.tl.read_gef function, generating spatial transcriptomic data with single-cell resolution for human fetal brain tissues. The entire pipeline can be accessed at https://stereopy.readthedocs.io/en/latest/Tutorials/Cell_Segmentation.html

### Spatial Transcriptomics clustering by FuseMap at single cell resolution

The FuseMap^81^ model was used to cluster ST data. Each spatial transcriptomics sample’s unique characteristics were modeled using dedicated autoencoders and graph autoencoders, forming a sample-specific probabilistic generative model. During the model training process, 15% of the cells were hold out as the validation set. Training was designed to early stop if there was no improvement in the validation loss for consecutive epochs. The training procedure included a pretraining phase, a weighted adversarial alignment as employed in previous research, and a final training phase that utilized cell-specific weights to ensure effective integration for cases when cell type compositions vary intrinsically. To further identify finer brain regions, Leiden^86^ clustering was applied to the spatial embedding of all spatial transcriptomics samples. Joint clustering was performed using the function scanpy.tl.leiden at a resolution 1.

### Mapping of snRNA data to ST data using SpatialID

We employed Spatial-ID^87^, an effective and reliable cell type annotation algorithm, to annotate cells within Stereo-seq data with the cell types clustered according to snRNA analyses to exhibit the spatial distribution of various cell types. Initially, we calculated the intersecting set of genes present in both the snRNA and Stereo-seq data. These genes were then used as input features during the training of a Deep Neural Network (DNN) model. Leveraging class-weight Cross Entropy Loss, the DNN model was trained over 30 epochs at a learning rate of 3e-4. This proficiently trained DNN model harbored the mapping information from gene profiles to predefined cell types. It was subsequently applied to cells in the Stereo-seq data to establish the initial probabilities of different cell types. Following this, the spatial neighborhood details of the Stereo-seq cells were encapsulated into an adjacency matrix. This matrix was weighted by normalized Euclidean distances. We then utilized a Graph Convolution Network (GCN) composed of two auto-encoders and a classifier block. This network integrated the gene profiles and spatial neighborhood information of the cells in spatial transcriptomic data to generate the final probabilities of cell types. The two auto-encoders received gene profiles and the adjacency matrix respectively, to create latent cell representations. Meanwhile, the classifier block established a connection from these representations to the initial cell type probabilities predicted by the DNN model. Training of these elements was achieved through self-supervised and classification losses using an adversarial learning strategy over a course of 200 epochs. The cell type that showed the highest value in the final probabilities was designated as the type of a cell. Spatial-ID utilized in this study was implemented using PyTorch (v1.13.1) and PyTorch Geometric (v2.1.0) in Python (v3.8.10).

### Cortical cell distribution analysis and visualization

For each ST data, the cell number of different cell types that mapped to different layers using SpatialID was quantified, and R package scatterpie (v0.1.8) was used to generate pie/ stacked bar chart showing the distribution of different cell types in each layer in the whole cortex or gyrus /sulcus groups. Differential abundance (gyrus vs. sulcus) was tested by Fisher’s exact test (FDR <0.1) and plotted as bubble maps (ggplot2 v3.4.2; size= -log10[FDR]).

### HotSpot gene module calculation

To identify gene enrichment modules in spatial transcriptomics data, a spatial adjacency matrix was constructed using positional data. Then, HotSpot^33^ was used to detect distinct gene modules. Finally, the significant gene module was obtained by applying an FDR threshold of <0.05.

### Gene module visualization

Gene module scores were calculated using Seurat AddModuleScore function, and visualized in ST data using SpatialPlot function.

### SpaTrack infer cell trajectory in ST data

SpaTrack^88^ was used to infer cell trajectory in ST data (n_neigh_pos = 100). The trajectory was then generated by using matplotlib.pyplot.streamplot function in Python.

### Ridge plot of genes distribution pattern

The tissue was rotated to make x axis align with dorsal-ventral axis. Then the cells expressed *TAC1* and/or *PENK* were quantified along x axis, and the density ridge plot was generated respectively.

### Calculate spatial variable genes (SVGs)

The region of interest was extracted using Spateo package as previously described. The spatial variable genes were calculated using Spateo package Moran I algorithm. The genes were filtered with Moran I score > 0.15 and p_val<0.05. Tope SVGs were plotted respectively.

### Gaussian blur and label transfer to extract patch and matrix

Selected SVGs were used to calculate the expression level on ST data. Gaussian blur was then used to extract patch and matrix structure from ST data. The annotation results from bin100 data were transferred to cellbin data. Specifically, a square region (side length: 100 arbitrary spatial units) was defined around the centroid coordinates of each bin100 point. All cellbin coordinates falling within this region were programmatically assigned the annotation corresponding to the central bin100 point, thereby establishing a spatially resolved annotation mapping between the two coordinate systems.

### Venn Diagram

DEGs for patch and matrix compartment were identified using the rank_gene_groups function in Scanpy. Genes exhibiting a log2 fold change (logFC) exceeding 0.5 were then intersected with those previously reported for patch and matrix regions in the referenced study^47^, respectively, and the overlaps were visualized using Venn diagrams.

### Polynomial regression

The Spateo Moran’s I algorithm was employed to identify spatially variable genes (SVGs) within individual ST section datasets. Significant genes were subsequently filtered by retaining those for which Moran’s I values in any three consecutive gestational weeks exceeded their gene-specific mean Moran’s I values computed across all gestational weeks. The polynomial regression analysis was then used to model the expression trends (ascending or descending) of genes across discrete gestational time points, based on z-score normalized and scaled gene expression profiles.

### Quantification and statistical analysis

This study did not use statistical methods to determine the sample size. The experiments were not randomized and the investigators were not blinded to the allocation during both the experiments and outcome assessment.

To assess region-specific differences in cellular abundance between gyrus and sulcus compartments, Chi-square tests were performed on integer-normalized cell type proportions (counts converted to whole-number percentages) with the chi2_contingency function in SciPy (v1.11.0). This normalization accounted for disparities in total cell counts across anatomical regions. For stereo-seq data of the gyrus and sulcus, following subregion annotation, differential expression analysis was performed using Wilcoxon Rank tests followed by Bonferroni correction for multiple tests. Statistical significance was set at p < 0.05, except for stereo-seq data which was set at padj < 0.01.

## Declaration of generative AI and AI-assisted technologies in the writing process

During the preparation of this paper, the authors used Copilot on STOmics Cloud (https://cloud.stomics.tech) to polish language. After using this tool, the authors reviewed and edited the content as needed and take full responsibility for the content of the publication.

## Data availability

- The raw sequence data have been deposited to CNGB Nucleotide Sequence Archive, and temporary access links can be requested from the lead contact. The data that support the findings of this study have been deposited into CNGB Sequence Archive (CNSA)^89^ of China National GeneBank DataBase (CNGBdb)^90^ with accession number CNP0005272 (http://db.cngb.org/cnsa/project/CNP0005272_8c0dc42f/reviewlink/).
- This paper does not provide the original code, the programs and packages used in the analysis are listed in the key resources table.
- Further details necessary for reanalyzing the data presented in this paper can be obtained upon request from the lead contact.

## Acknowledgments

This work was supported by Sichuan Science and Technology Program (2024YFFK0072); National Key Research and Development Program of China (2022YFC3400400 to X.X.); National Key R&D Program of China (2021YFA0805100); the Frontiers Medical Center, Tianfu Jincheng Laboratory Foundation (TFJCPI20250038) and the STI 2030-Major Projects (2021ZD0200500). Project analysis was performed on the STOmics Cloud (https://cloud.stomics.tech). This paper is from the Mesoscopic Brain Mapping Consortium.

## Author contributions

L.Z, X.L and Yaoyao.Z designed the project; Yunjia.Z, Y.L, Xunan.S, Xiaobo.S, T.Z, Z.W, Z.Z, L.C, S.Q, J.Y, and F.H performed analysis; Investigation, Xue.X, C.S, C.L, M.L, F.W, Yaqi.L and J.L performed experiments and discussion; Yanxin.L, J.J., Xue.X, and Yaoyao.Z collected the samples; Yunjia.Z, Y.L, Xunan.S, and C.S wrote the draft,; Yunjia.Z, Y.L, Xunan.S, C.S, Q.Z, Z.L, S.Z, Z.D, and L.L reviewed and edited the manuscript; Yunjia.Z, L.Z, S.L, and Yaoyao.Z supervised the project; X.X, S.L, L.Z, and Yaoyao.Z provide funding support.

## Additional information

**Correspondence** and requests for materials should be addressed to Yaoyao Zhang.

---

**Fig.S1.**
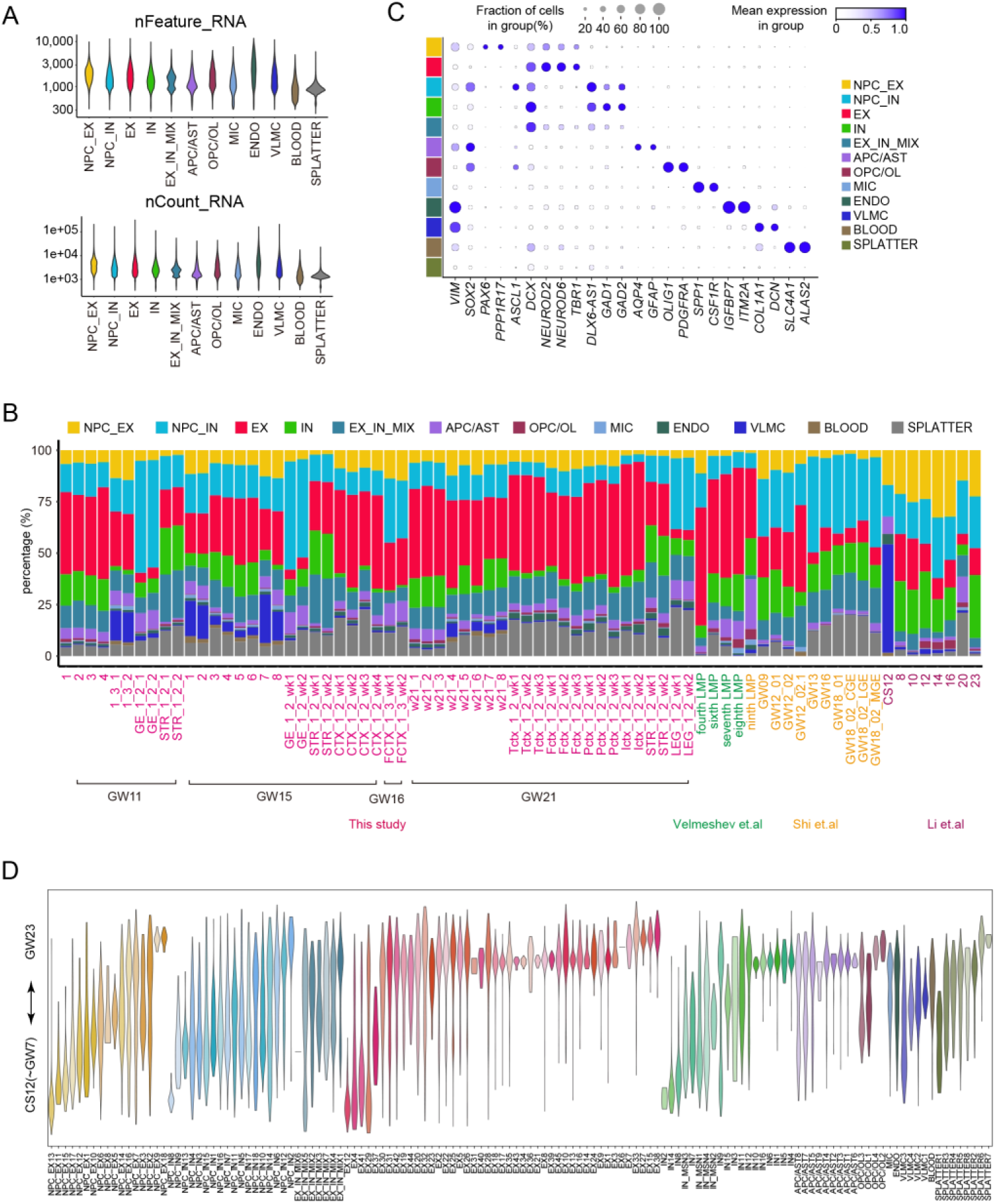
| Quality control, major cell types and subtypes of snRNA-seq data, related to Fig.1. **A.** Quality control of snRNA-seq clusters by counts and RNA features. **B.** Stack-bar plot showing the percentage of major cell types in each library and sample. **C**. Bubble plot showing the expression levels of the specific markers in each of the major cell types. Bubble size shows the percentage of cell expressed; scale bar shows the average expression level. **D.**MILO analysis showing the distribution of 134 (out of 136 cell subtypes, NPC_EX4 and IN13 were removed due to low MILO value) cell subtypes along developmental stages.

**Fig.S2.**
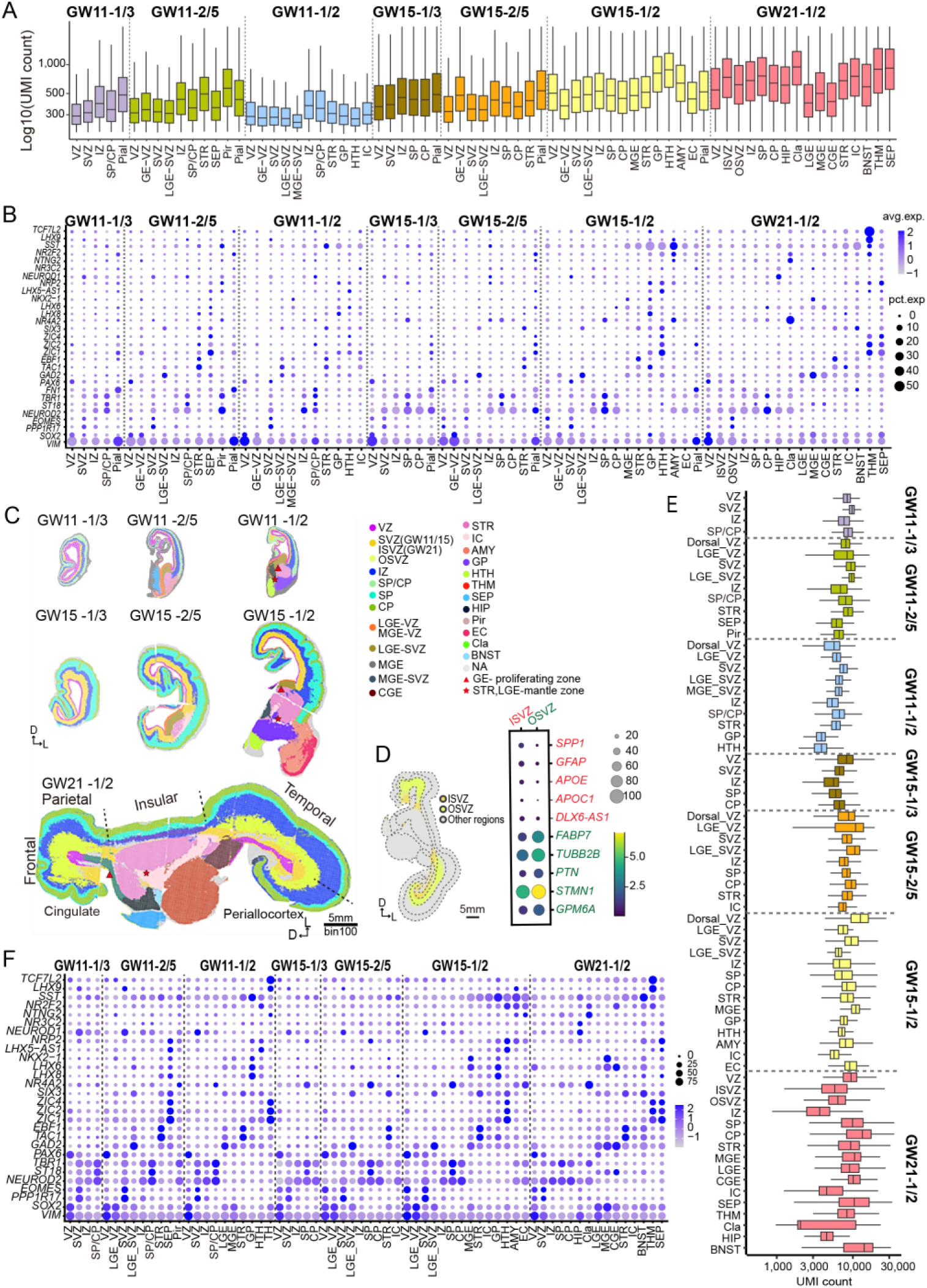
| Quality control, spatial region registration of stereo-seq data, related to Fig.1. **A.** Box plot showing the number of detected UMIs per brain region on spatial transcriptomic data, clustered by FuseMap at single cell resolution. **B.** Bubble plot showing the expression levels of the specific markers in each of the sub-brain regions, clustered by FuseMap at single cell resolution. Bubble size shows the percentage of cell expressed; scale bar shows the average expression level. **C.** Spatial clustering of stereo-seq brain regions by SpaGCN at bin100 resolution. Brain regions are colored by the annotations. VZ: ventricle zone; SVZ: sub-ventricle zone; ISVZ: inner sub-ventricle zone; OSVZ: outer sub-ventricle zone; IZ: intermediate zone; SP: subplate; CP: cortical plate; LGE: lateral ganglionic eminence; MGE: medial ganglionic eminence; CGE: caudal ganglionic eminence; STR: striatum (caudate and putamen); IC: internal capsule; AMY: amygdala; GP: globus pallidus; HTH: hypothalamus; THM: thalamus; SEP: septum; HIP: hippocampus; Pir: piriform cortex; EC: entorhinal cortex; Cla: claustrum; BNST: bed nucleus of striata terminal; NA: not available. D: dorsal; L: lateral. Scales: 5mm. **D.** Identification of ISVZ and OSVZ regions as well as key DEGs expression. Spatial subclustering of stereo-seq SVZ regions by SpaGCN at bin100 resolution. SVZ subregions are colored by the annotations. ISVZ: inner subventricular zone; OSVZ: outer subventricular zone. D: dorsal; L: lateral. Scales: 5 mm (Left). Bubble plot showing the expression levels of the top 5 DEGs enrichment in ISVZ and OSVZ. Bubble size shows the percentage of square bins expressed; the scale bar shows the average expression level (Right). **E.** Box plot showing the number of detected UMIs per brain region on spatial transcriptomic data, clustered by SpaGCN at bin100 resolution. **F.** Bubble plot showing the expression levels of the specific markers in each of the sub-brain regions, clustered by SpaGCN at bin100 resolution. Bubble size shows the percentage of cell expressed; scale bar shows the average expression level.

**Fig.S3.**
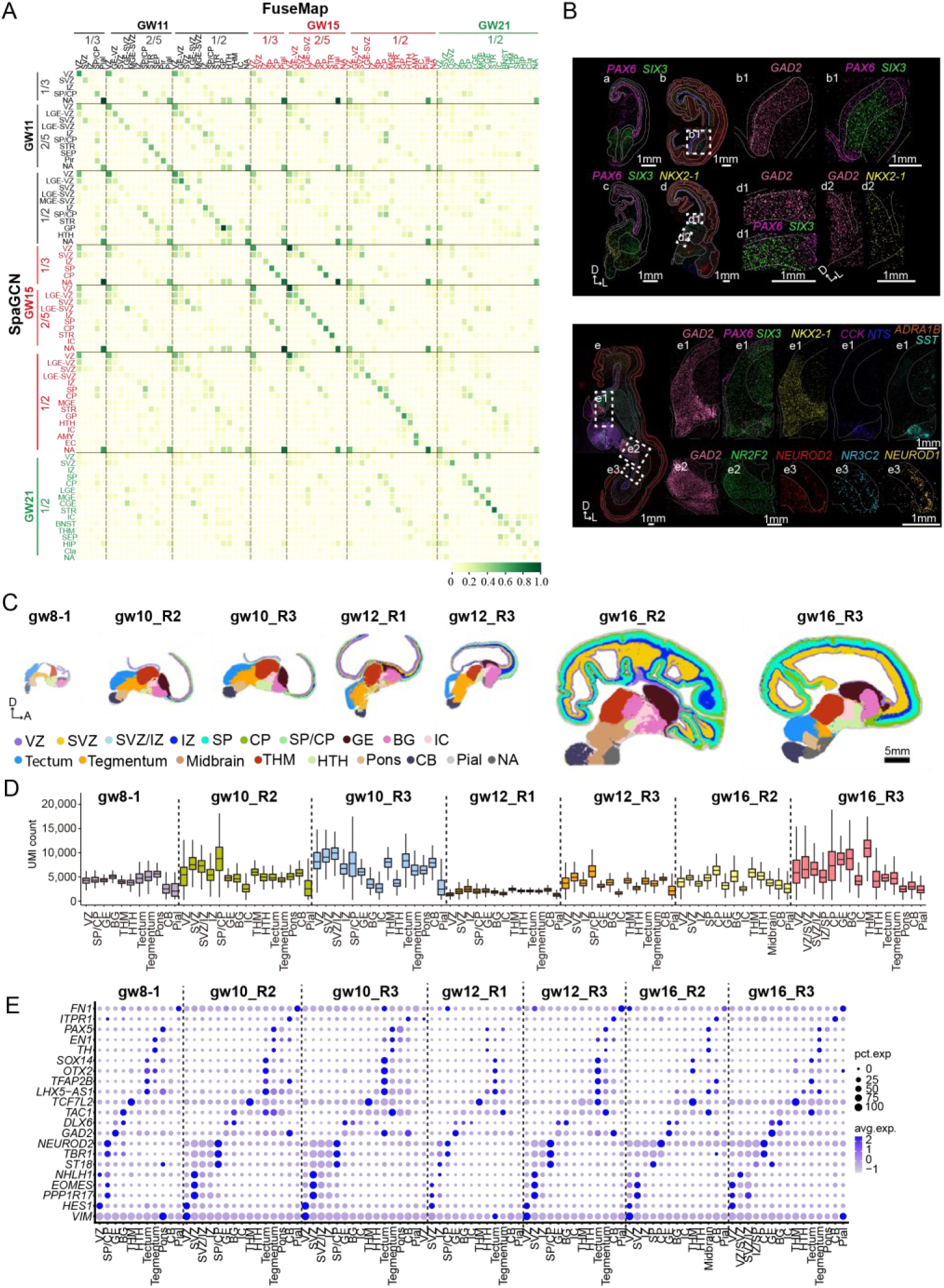
| Quality control, spatial region registration of stereo-seq data, related to Fig.1. **A.** Heatmap illustrating the comparison of spatially defined brain subregions clustered by FuseMap and SpaGCN. **B.** Spatial visualization of specific brain region markers. White solid lines represent the borders of subregions. HIP: *NEUROD1*, *NR3C2*; LGE-proliferating zone: *PAX6*, *SIX3*; MGE-proliferating zone: *NKX2-1*, CGE-proliferating zone: *NR2F2*; BNST: *CCK*, *NTS*, *SST*, *ADRA1B*. White dash lines represent the higher-magnification view of selected area. D: dorsal; L: lateral. Scales: 1mm. **C.** Spatial clustering of stereo-seq brain regions from Li’s data by SpaGCN at bin100 resolution. Brain regions are colored by the annotations. VZ: ventricle zone; SVZ: sub-ventricle zone; IZ: intermediate zone; SP: subplate; CP: cortical plate; GE: ganglionic eminence; BG: basal ganglia; IC: internal capsule; THM: thalamus; HTH: hypothalamus; CB: cerebellum; NA: not available. D: dorsal; A: anterior. Scales: 5mm. **D.** Box plot showing the number of detected UMIs per brain region on spatial transcriptomic data from Li et. al, clustered by SpaGCN at bin100 resolution. **E.** Bubble plot showing the expression levels of the specific markers in each of the brain regions from Li et. al, clustered by SpaGCN at bin100 resolution. Bubble size shows the percentage of cell expressed; scale bar shows the average expression level.

**Fig.S4.**
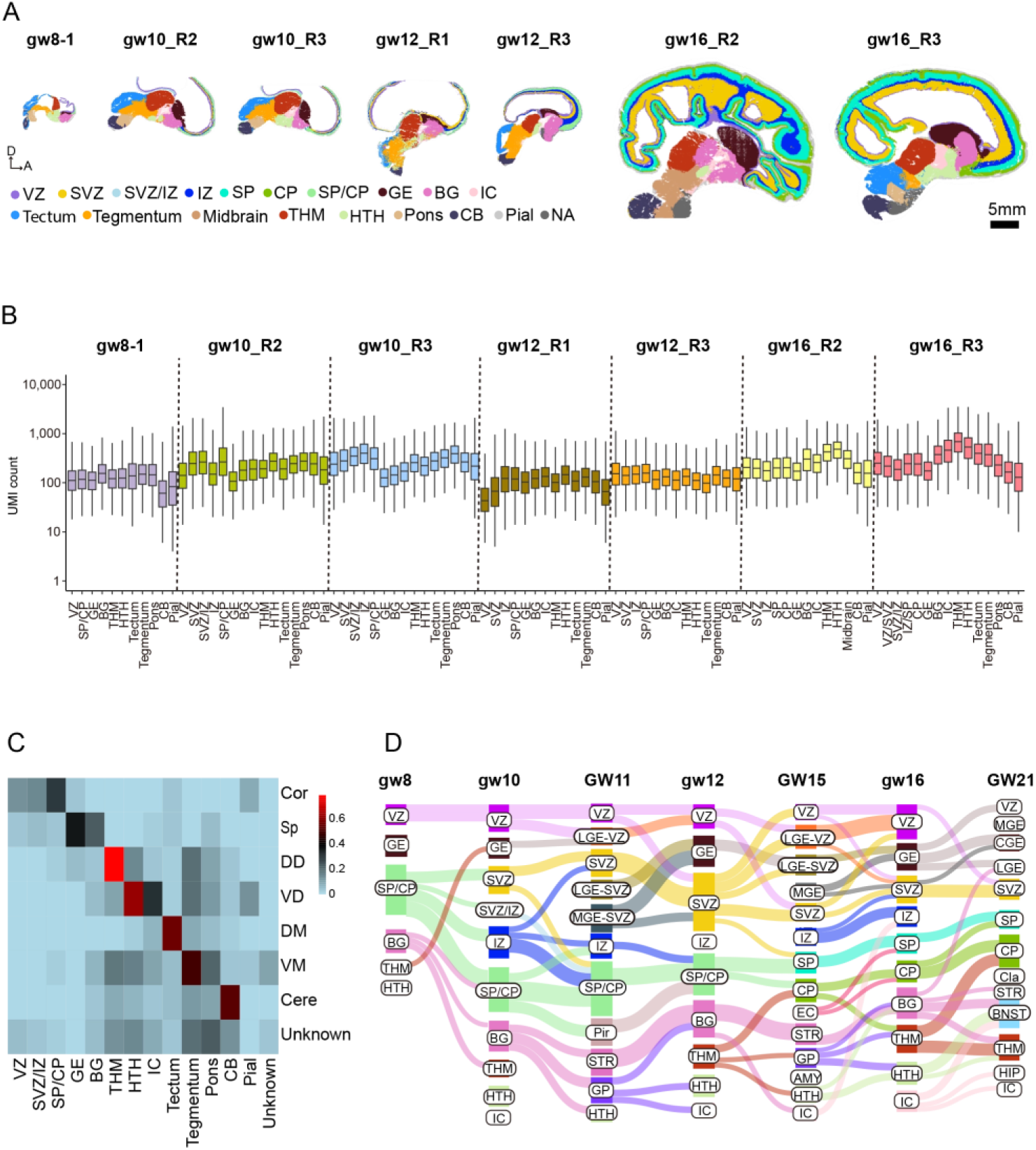
| Spatial region registration and comparison of stereo-seq data, related to Fig.1. **A.** Spatial annotation of stereo-seq brain regions from Li’s data by SpaGCN at single cell resolution. Brain regions are colored by the annotations. VZ: ventricle zone; SVZ: sub-ventricle zone; IZ: intermediate zone; SP: subplate; CP: cortical plate; GE: ganglionic eminence; BG: basal ganglia; IC: internal capsule; THM: thalamus; HTH: hypothalamus; CB: cerebellum; NA: not available. D: dorsal; A: anterior. Scales: 5mm. **B.** Box plot showing the number of detected UMIs per brain region on spatial transcriptomic data from Li et. al, annotated by SpaGCN at single cell resolution. **C.** Heatmap illustrating the comparison of spatially defined brain subregions clustered by SpaGCN with anatomically defined brain regions from Li et. al. **D.** Sanky plot showing the correlation of brain regions across developmental stages. GW: samples from this study; gw: published data by Li et al.

**Fig.S5.**
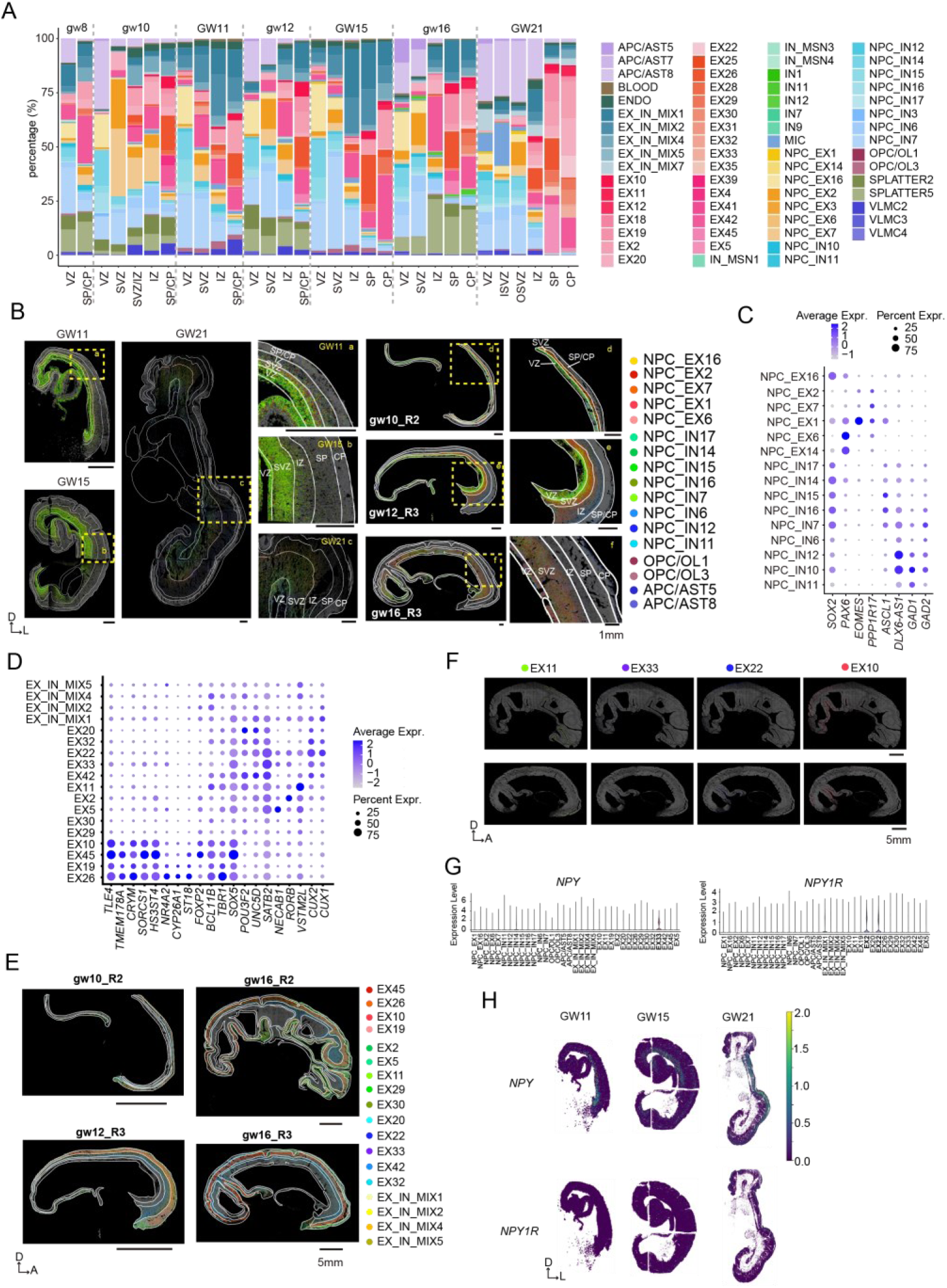
| The dynamic distribution of cells in cortical expansion and arealization, related to Fig.2. **A.** Stack-bar plot showing the percentage of cells in each spatial cortical layer during development. **B.** Spatial plot showing 13 NPC cell types and glial cells at single cell resolution across different developmental stages. White solid lines represent the borders of sublayers. Yellow dotted boxes represent the zoom in region. D: dorsal; L: lateral. Scales:1mm. **C.** Bubble plot showing the expression levels of the specific markers in NPC clusters from snRNA data. Bubble size shows the percentage of cell expressed; scale bar shows the average expression level. **D.** Bubble plot showing the expression levels of the deep layer and upper layer markers in EX clusters from snRNA data. Bubble size shows the percentage of cell expressed; scale bar shows the average expression level. **E.** Spatial plot showing 14 excitatory cell types and 4 EX_IN_MIX cells at single cell resolution. White solid lines represent the borders of sublayers. D: dorsal; A: anterior. Scales:5mm. **F.** Spatial plot showing the distribution of 4 excitatory cell types in different cortical lobes at gw16. D: dorsal; A: anterior. Scales:5mm. **G.** Violin plot showing the expression level of the ligand NPY and the receptor NPY1R in snRNA data. **H.** Spatial visualization of the expression of the ligand NPY and the receptor NPY1R at GW21. D: dorsal; L: lateral.

**Fig.S6.**
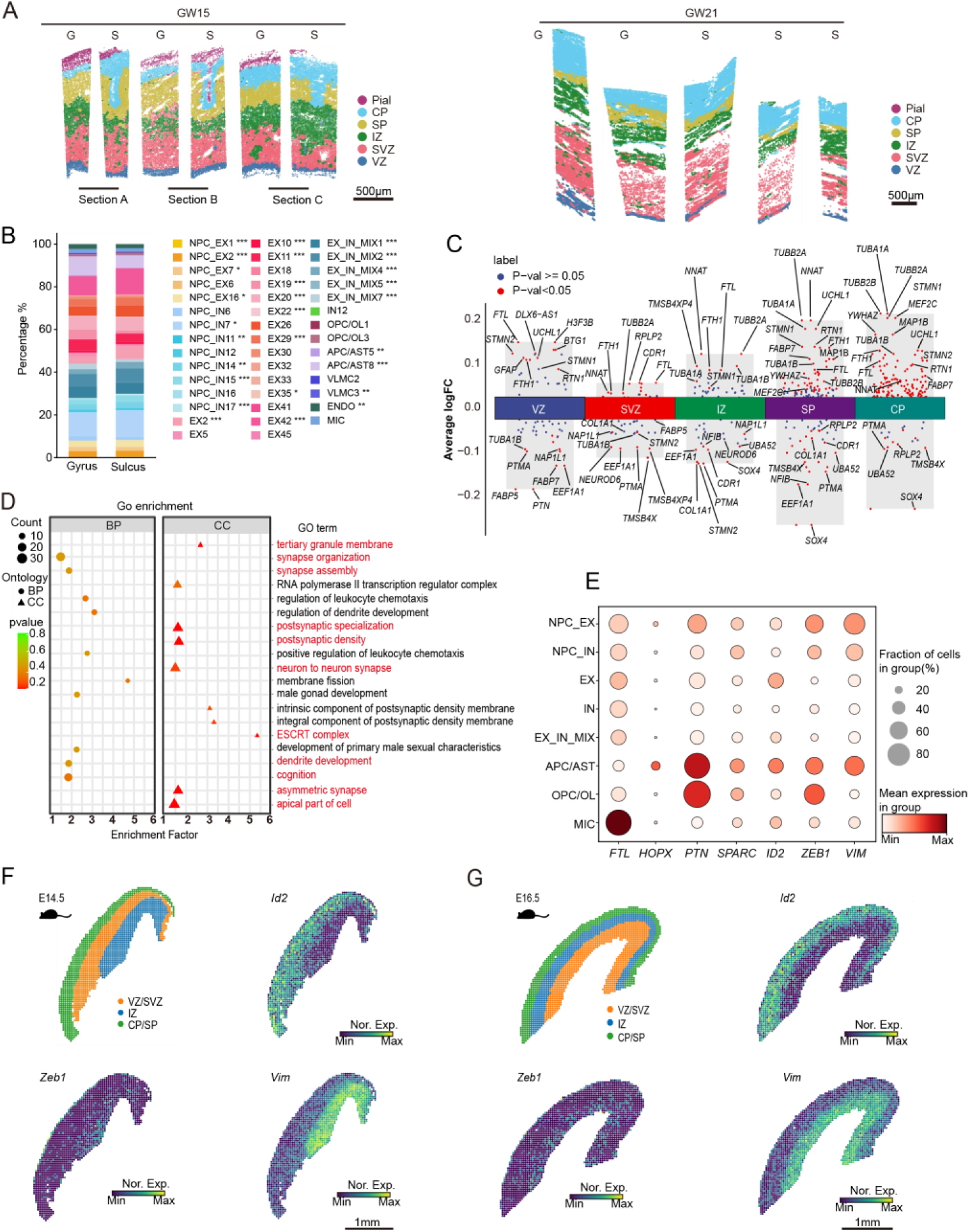
| The occupied region, transcriptomic and the cellular heterogeneity in developing gyrus and sulcus, related to Fig.3. **A.** Spatial clustering of lassoed gyrus and adjacent sulcus region stereo-seq coronal sections at GW15 and GW21 by FuseMap. Subregions are colored by the annotations: VZ (blue), SVZ (pink), IZ (green), SP (yellow), CP (light blue), Pial(purple). Scales: 500μm. **B.** Stack-bar plots showing the percentage of cell subtypes between gyrus and sulcus. (Chi-squared test with Benjamini-Hochberg correction, *p<0.05, **p<0.01, ***p<0.001). **C.** Volcano plots illustrate gyrus-sulcus comparisons in five neurodevelopmental subregions: VZ, SVZ, IZ, SP, and CP. Significantly DEGs are defined by Benjamini-Hochberg adjusted *p* < 0.05 (two-tailed Wald test). Color encoding: red (p < 0.05), navy blue (non-significant, p ≥ 0.05). **D.** GO terms for genes upregulated in gyrus (p-value: Fisher’s exact test, one-sided). **E.** Bubble plot illustrates the expression levels of seven DEGs (*FTL, HOPX, PTN, SPARC, ID2, ZEB1, VIM*) demonstrating significant gyral enrichment across snRNA-seq defined cortical cell clusters. Bubble size shows the percentage of cell expressed; scale bar shows the average expression level. **F.** Spatial expression of *Id2*, *Zeb1* and *Vim* in mouse E14.5 cortex sagittal section (MOSTA data). Scales: 1mm. **G.** Spatial expression of *Id2*, *Zeb1* and *Vim* in mouse E16.5 cortex sagittal section (MOSTA data). Scales: 1mm.

**Fig.S7.**
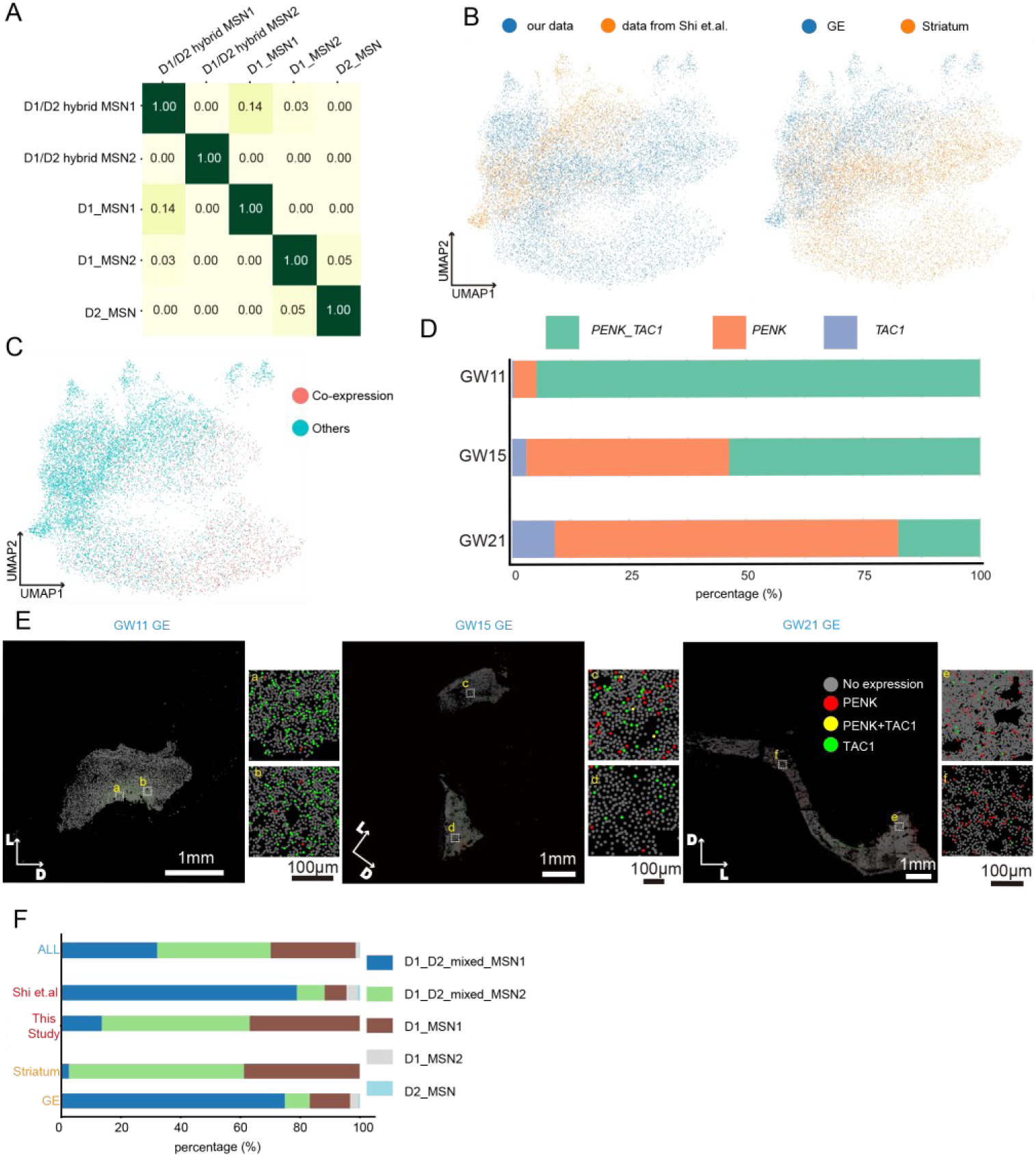
| Validation and comparative analysis of MSN subtypes, related to Fig.4. **A.** Jaccard similarity index quantifying overlap between MSN subtype classifications across independent clustering iterations. **B.** UMAP projection showing different data source and tissue regions. Left: UMAP projection integrating snRNA-seq data from this study and a reference dataset (Shi et al.). Right: UMAP stratification of MSNs by tissue origin (GE vs. striatum). **C.** UMAP visualization illustrating co-expression of *TAC1* and *PENK* across MSN subtypes in snRNA-seq data. **D.** Stack bar plot showing the distribution of *TAC1* and *PENK* as well as co-expression in different gestational weeks in striatum. **E.** Spatial transcriptomics colormap showing *TAC1* and *PENK* co-expression within the GE. Scales: white = 1 mm, black = 100 μm; inset shows higher-magnification view. **F.** Bar plot comparing the proportional representation of MSN clusters across integrated snRNA-seq dataset.

**Fig.S8.**
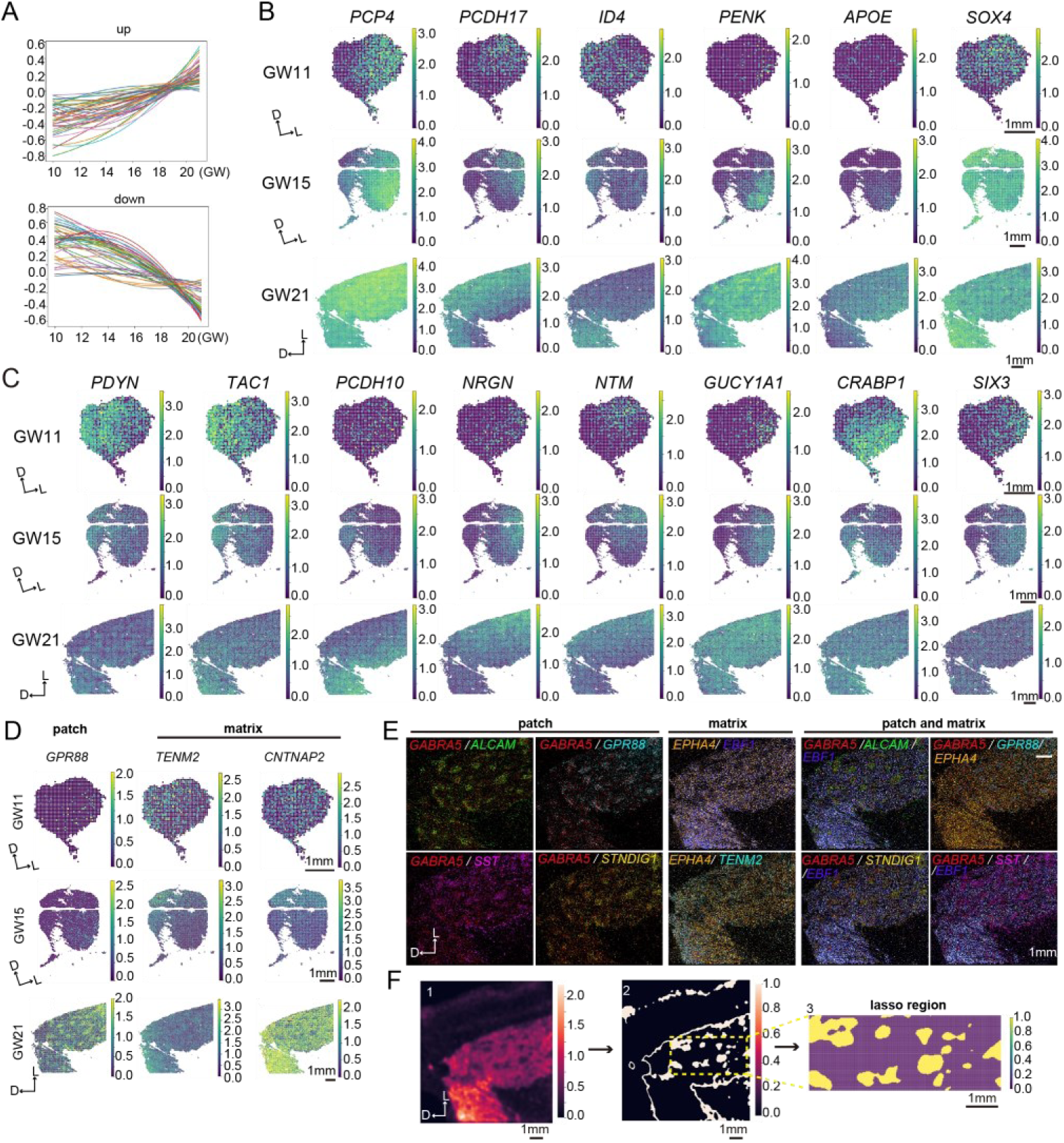
| Spatial transcriptomic identification of patch and matrix compartments, related to Fig.5. **A.** Polynomial regression showing the up- or down-expression trend of genes across discrete gestational time points. The genes were z-score normalized and scaled. **B.** Spatial gene expression plot of significant genes selected from Fig S8A. Scales:1mm. **C.** Spatial expression of dorsal and ventral enriched genes at GW11, GW15, GW21. D: dorsal; L: lateral. Scales:1mm. **D.** Spatial expression of patch and matrix enriched genes at GW11, GW15, GW21. D: dorsal; L: lateral. Scales:1mm. **E.** Colocalization of novel and well-known patch/matrix genes. D: dorsal; L: lateral. Scales:1mm. **F.** Gaussian blur identifying regions of patch and matrix by SVGs at GW21. D: dorsal; L: lateral. Scales:1mm.

**Fig.S9.**
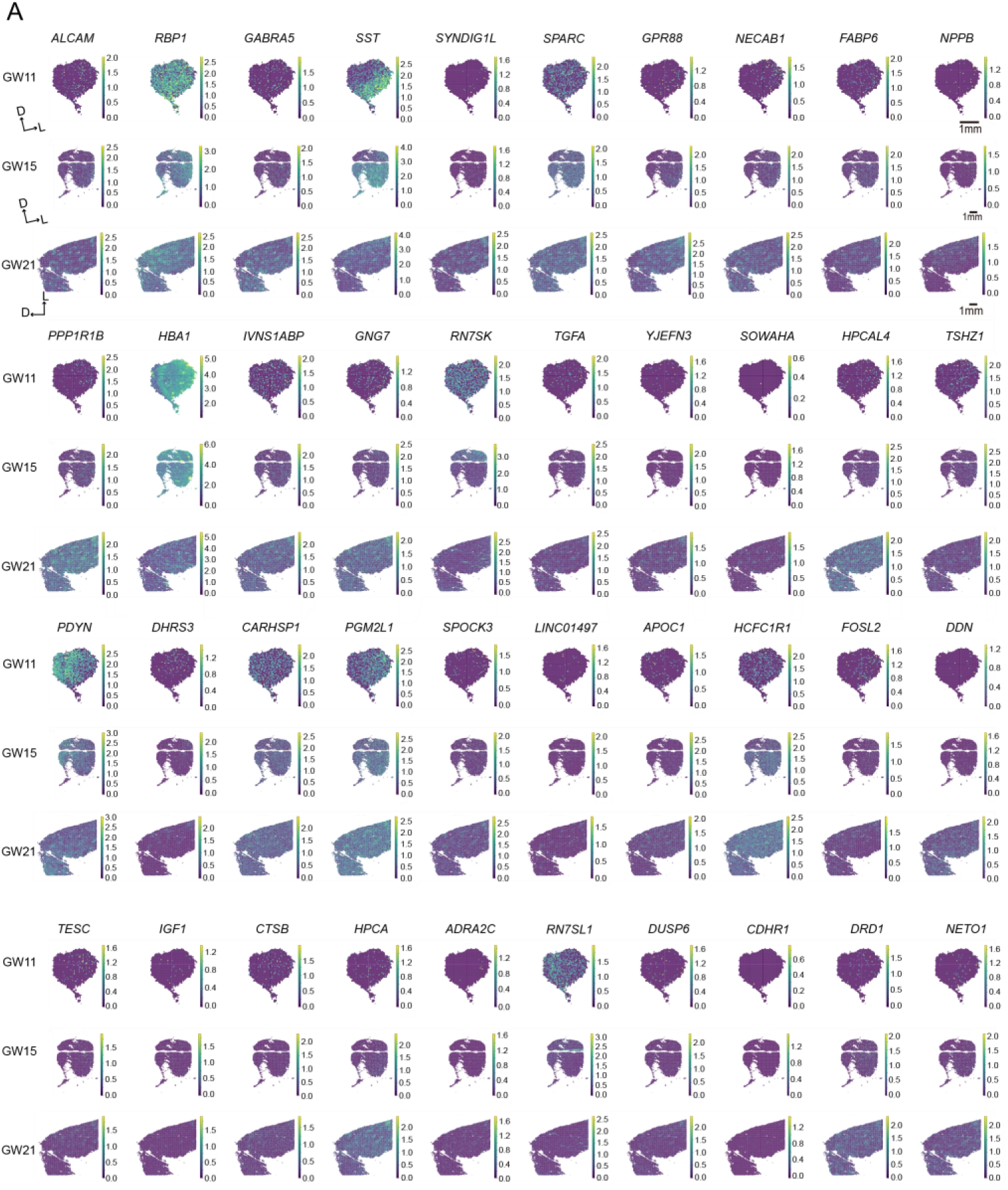
| Spatial visualization of top 40 patch DEGs, related to Fig.5. **A.** Spatial visualization of top 40 patch DEGs. D: dorsal; L: lateral. Scales:1mm.

**Fig.S10.**
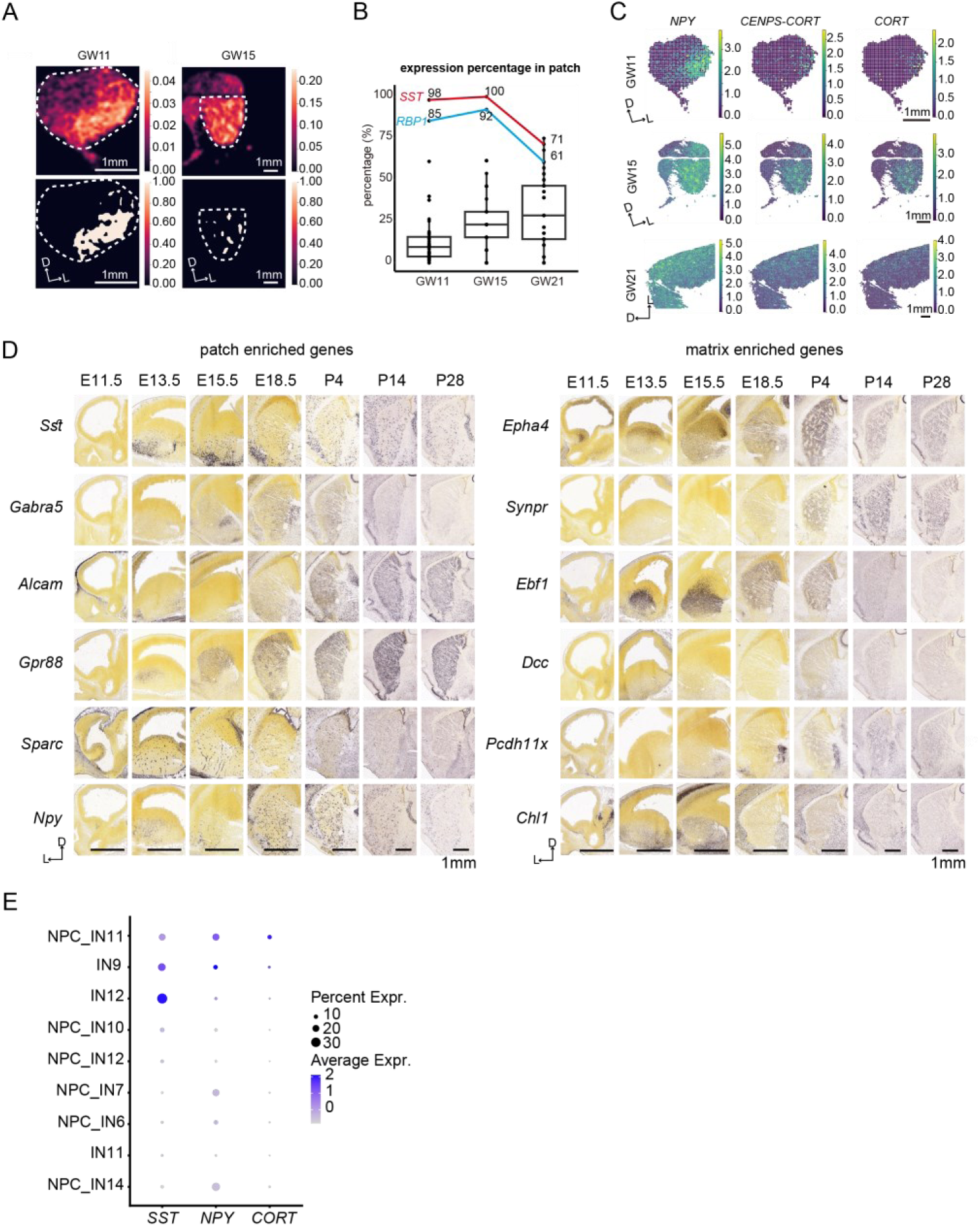
| Spatial analysis of patch and matrix DEGs, related to Fig.5. **A.** Gaussian blur identifying regions of patch and matrix by SVGs at GW11 and GW15. D: dorsal; L: lateral. Scales:1mm. **B.** The fraction of cells expressing patch marker genes at GW11, GW15 and GW21. Each dot represents a patch DEG calculated. The red and blue lines show two representative high expression genes in patch. **C.** Spatial visualization of neuropeptide genes *NPY, CENPS-CORT* and *CORT* at GW11, GW15 and GW21. D: dorsal; L: lateral. Scales: 1mm. **D.** Expression of patch and matrix enriched top genes in ISH data of mouse brain at E11.5, E13.5, E15.5, E18.5, P4, P14 and P28 from Allen Brain Atlas. Scales: 1mm. **E.** Dot plot showing the expression of *SST, NPY* and *CORT* in striatum interneurons from snRNA data.

## supplemental Excel table titles and legends

Table S1. Hotspot gene module, related to Figure 2

Table S2. Cell proportions of gyri and sulci in each sublayer, related to Figure 3

Table S3. DGEs of gyri and sulci in each sublayer, related to Figure 3

Table S4. DEGs and GO pathways of patch and matrix at GW21, related to Figure 5

